# *Rhodococcus parequi* sp. nov., a new species isolated from equine farm soil closely related to the pathogen *Rhodococcus equi*

**DOI:** 10.1101/2024.12.09.627583

**Authors:** José A. Vázquez-Boland, Jorge Val-Calvo, Fabien Duquesne, Francesca Decorosi, Carlo Viti, Sandrine Petry, Mariela Scortti

## Abstract

We present the description of the new species, *Rhodococcus* (*Prescottella*) *parequi*, found during phylogenomic investigations of a global collection of strains identified as *Rhodococcus* (*Prescottella*) *equi*. Strain PAM 2766 was isolated from horse-breeding farm soil in Normandy, France, and was indistinguishable from *R. equi* based on the usual identification tests. Whole-genome phylogenetic analyses located PAM 2766 in the same *Rhodococcus* sublineage as *R. equi*, together with *Rhodococcus agglutinans*, *Rhodococcus defluvii*, *Rhodococcus soli*, *Rhodococcus subtropicus*, *Rhodococcus spongiicola* and *Rhodococcus xishaensis*. PAM 2766 is most closely related to, but sufficiently distinct from *R. equi* DSM 20307^T^ to be considered as a separate species. Average Nucleotide Identity (ANI) and Average Amino Acid Identity (AAI) values are 88.60% and 92.35, respectively, well below the species cutoff. The PAM 2766 draft genome is ∼5.3 Mb in size with 68.98% G+C mol content. PAM 2766^T^ is aerobic, non-motile, and produces smooth creamy to buff-coloured colonies very similar to those of *R. equi*. It phenotypically differs from the latter by the ability to grow at 5°C, a strongly positive urease test at 24 h, and specificities in the carbon and nitrogen source utilization profile as determined by phenotype microarray screens. Our data indicate that PAM 2766 belongs to a novel species, for which the name *Rhodococcus parequi* sp. nov. is proposed. *R. parequi* was avirulent in macrophage infection assays and is assumed to be non-pathogenic. The type strain is PAM 2766^T^ (=CETC 30995^T^ = NCTC 14987^T^).

## INTRODUCTION

The bacterial genus *Rhodococcus* Zopf 1891 (Approved Lists 1980) [1, 2], within the *Nocardiaceae* family of the *Mycobacteriales*, comprises a large group of irregular Gram-positive, aerobic coccobacilli with more than half a hundred recognized species and an ever-growing number of unclassified isolates (LPSN, https://www.bacterio.net [3]). The rhodococci are ubiquitous saprophytes that can be isolated from a wide diversity of habitats including soil, waters, marine sediments and extreme environments such as deep sea, caves and xenobiotic-contaminated sites [4–6]. One member of the genus, *Rhodococcus equi* commonly found in soil, is a major equine pathogen and human opportunistic pathogen [7, 8]. This study reports the isolation of a novel *Rhodococcus* species that is closely related to *R. equi*, with potential implications in the identification of this pathogen and in studies on its ecology, epidemiology and environmental distribution. Here we use the name *R. equi* instead of the synonym *Prescottella equi* (Magnusson 1923) Sangal *et al*. 2022 [9], and the circumscription of the genus *Rhodococcus* Zopf 1891 as emended by Val-Calvo and Vázquez-Boland 2023 [10]. This emendation was based on a recent study that examined the *Mycobacteriales* taxonomy using a network analysis-aided, context-uniform phylogenomic approach for non-subjective genus demarcation [11].

### STRAIN ISOLATION

Strain PAM 2766 was isolated from a culture kept in the Vazquez-Boland’s laboratory collection as *R. equi* strain PAM 1352. The PAM 1352 culture was originally obtained in 1999 from a soil sample from a stud farm in Normandy, France. PAM 1352 was classified as *R. equi* based on colony morphology, API Coryne (BioMériux) biochemical profiling, detection of cholesterol oxidase activity by the CAMP-like synergistic haemolysis test with *Listeria ivanovii* (a strong sphingomyelinease C producer) [12, 13], and a positive PCR for the *R. equi* cholesterol oxidase gene *choE* (considered as a species-specific identification marker [14–17]). In a previous study, PAM 1352 was found to lack the *R. equi* virulence plasmid by TRAVAP typing, a PCR-based method to differentiate the three *R. equi* virulence plasmid types [16, 18]. During a population phylogenomics study of the *R. equi* species, WGS analysis of PAM 1352 suggested the culture was contaminated with a closely related taxonomically unidentified microorganism. Meticulous re-isolation followed by WGS revealed that PAM 1352 was a mixed culture of two different bacteria with very similar colonial morphologies: one corresponded to *R. equi*, the other to a putative new species. The latter was assigned PAM number 2766 and deposited at the NCTC repository under number 14987 and CECT repository under number 30995 (henceforth PAM 2766^T^, NCTC 14987^T^ and CECT 30995^T^, respectively).

### GENOMIC CHARACTERIZATION

Genomic DNA of PAM 2766^T^ was isolated (GenEluteTM Bacterial Genomic DNA, Sigma) and sequenced by Illumina with 150-bp paired-end reads and minimum 150× coverage (Novogen). The raw sequencing data were processed using FastQC (https://www.bioinformatics.babraham.ac.uk/projects/fastqc/) and Trimmomatic [19] (settings: Leading 3, Trailing 3, Slidingwindow 4:15, Minlen 36) to remove low-quality reads and adapter sequences, then assembled using SPAdes v3.15.2 [20] (option isolate active, k-mer length and read coverage set to auto). The draft PAM 2766^T^ genome consisted of 16 contigs of >500 bps with a total length of 5,295,699 bps and a G+C content of 68.98 mol%. Actual sequence coverage depth was 278× and contig scores of N50 = 611,300 and L50 = 4 determined using QUAST v5.1.0rc1 [21] were consistent with a quality assembly for taxonomic purposes. The PAM 2766^T^ draft genome was annotated using NCBI Prokaryotic Genome Annotation Pipeline (PGAP) [2023-10-03.build7061] [22].

When aligning in QUAST the PAM 2766^T^ genome contigs (unfractionated) to the 103S reference genome (the first complete and only manually curated genome for *R. equi*; GenBank accession no. FN563149.1) [23], several metrics suggested that it deviated from a typical *R. equi* genome assembly. For example, the Genome Fraction % (percentage of aligned contig bases in the reference genome) and Total Aligned Length (number of aligned contig bases in the assembly) were 1.04% and 53,807 for PAM 2766^T^ vs 95.23% and 4,804,699 for the *R. equi* DSM 20307^T^ genome, respectively. Similar diverging values were obtained with other random *R. equi* genome assemblies.

The genomic distinctiveness of PAM 2766^T^ was confirmed using two overall genomic relatedness indices (OGRIs) recognized as robust indicators for species demarcation, i.e. the Average Nucleotide Identity (ANI) and Average Amino Acid Identity (AAI) [22, 24–26]. The values for both indices between PAM 2766^T^ and *R. equi* DSM 20307^T^ were well below the <95% species definition cutoff (ANI = 88.60 ± 4.11; AAI = 92.35 ± 7.42). In silico DNA-DNA hybridization (dDDH) analyses performed using the genome-genome distance calculator 3.0 (GGDC) [27] also reported a low probability (1.4%) for a dDDH value greater than the species delimitation threshold of >70% [25]. Next in relatedness were the *Rhodococcus* species located within the same phylogenomic sublineage as *R. equi* (rhodococcal sublineage no. 2 [11]) (**Fig. 1**). ANI/AAI values ranged from 85.21/85.80 for *Rhodococcus agglutinans* to 82.00/81.32, 82.13/80.73 and 82.83/80.87 for *Rhodococcus xishaensis*, *Rhodococcus spongiicola* and *Rhodococcus subtropicus*, respectively (**Table 1**) *R. soli* DSM 46662^T^ was *de novo* sequenced in this study (GenBank accession no. JBDLNU000000000) as its genome information was not publicly available.

**Fig. 1.**
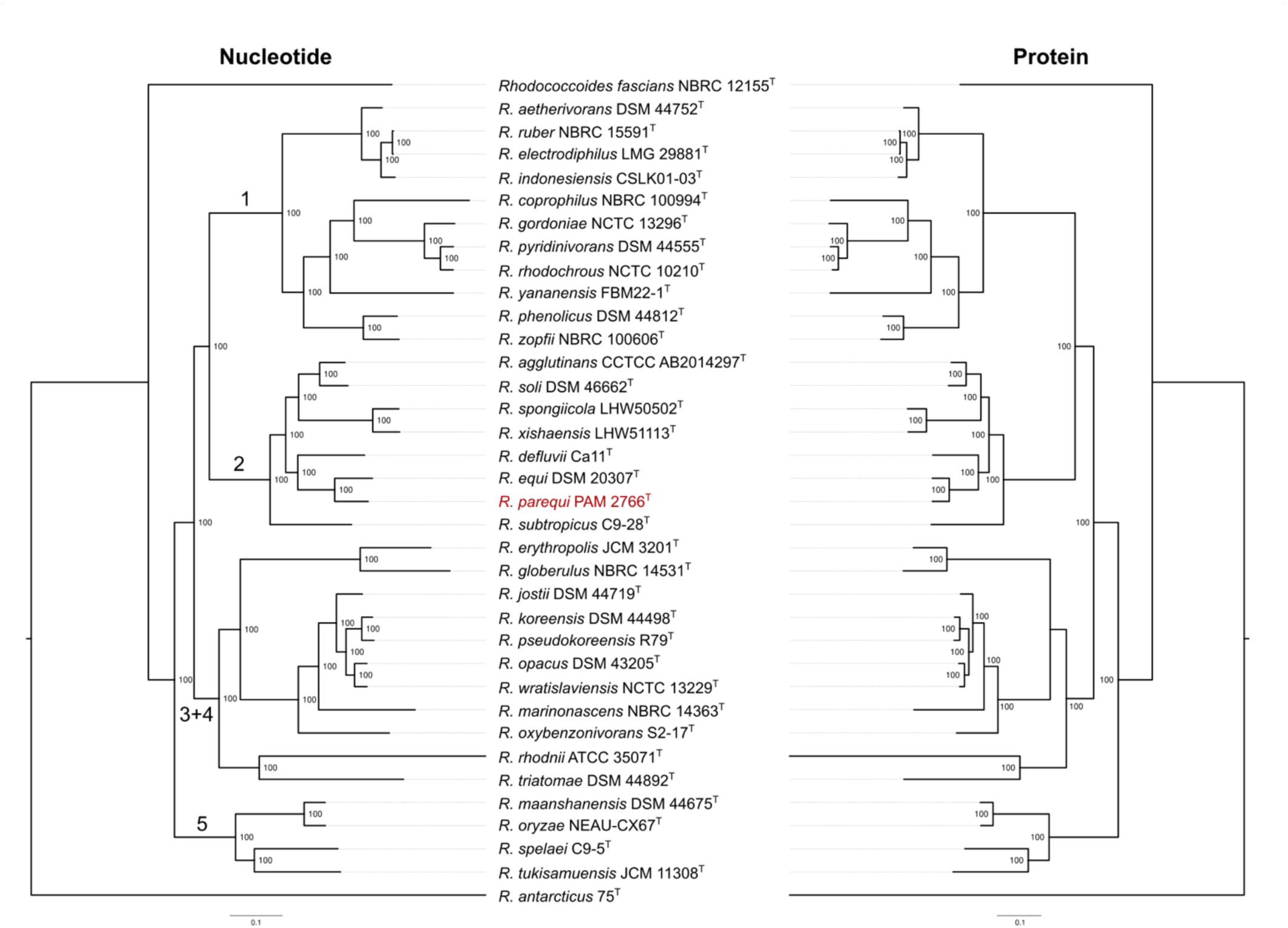
Core-genome ML trees based on concatenated sequence alignments of conserved genes (left, 409 gene markers) or proteins (right, 323 protein markers). The analysis includes the type strains of all *Rhodococcus* species with genome assemblies available at NCBI as of January 2024. *Rhodococcoides fascians* from an early diverging branch of the rhodococcal radiation [11] was used as an outgroup. The tree was midpoint rooted because of the uncertain position of *Rhodococcus antarcticus* 75^T^ within the genus *Rhodococcus*. Branch support is given for 1,000 ultrafast bootstrap replicates (IQtree UFBoot). The large case numbers on the nucleotide-based tree indicate the main *Rhodococcus* sublineages as defined in ref. [11]. The trees are entirely congruent with our previous ML phylogenies based on *Mycobacteriales* or *Nocardiaceae* core-genome alignments (the latter incorporating a larger strain representation than the type’s) [11]. The only exception is that *R. rhodnii* ATCC 33071^T^ and *R. triatomae* DSM 44892^T^ form here an independent branch at the root of the monophyletic line of descent containing sublineages 3 and 4 instead of being part of the sublineage 4 radiation (see ref. [11]).

**Table 1.** Genome relatedness index comparisons between *R. parequi* PAM 2766^T^ and most closely related species (*Rhodococcus* phylogenomic sublineage no. 2 [11]).

|  | 16S<sup>a</sup><br>% similarity | ANI<sup>b</sup> | AAI<sup>b</sup> | GGD<sup>c</sup><br>(% probability) |
| --- | --- | --- | --- | --- |
| *R. agglutinans* CFHS 0262<sup>T</sup> | 99.03 | 85.21 ± 5.19 | 85.80 ± 11.48 | 0.1522 (0.05) |
| *R. defluvii* Ca11<sup>T</sup> | 98.89 | 84.52 ± 5.13 | 85.31 ± 11.68 | 0.1565 (0.03) |
| *R. equi* DSM 20307<sup>T</sup> | 100<sup>d</sup> | 88.60 ± 4.11 | 92.35 ± 7.42 | 0.1079 (1.44) |
| *R. soli* DSM 46662<sup>T</sup> | 99.31 | 84.57 ± 5.08 | 84.95 ± 12.23 | 0.1558 (0.04) |
| *R. spongiicola* LHW50502<sup>T</sup> | 97.15 | 82.13 ± 4.65 | 80.73 ± 14.12 | 0.1808 (0.01) |
| *R. subtropicus* C9-28<sup>T</sup> | 98.68 | 82.83 ± 4.94 | 80.87 ± 13.30 | 0.1763 (0.01) |
| *R. xishaensis* LHW51113<sup>T</sup> | 97.15 | 82.00 ± 4.63 | 81.32 ± 13.48 | 0.1804 (0.01) |

<sup>a</sup> Determined using EzBioCloud 16S database [53], except for *R. equi* (see footnote d).
<sup>b</sup> Calculated using the scripts ani.rb and aai.rb from the “Enveomics collection” [54].
<sup>c</sup> Genome-to-genome distance calculated based on the recommended formula 2 (identities/HSP length). In parentheses, the probability of a DDH > 70% (logistic regression).
<sup>d</sup> Based on the 16 rDNA sequence of DSM 20307<sup>T</sup> genome assembly GenBank accession no. LWTX01000024 (LWTX01000000 – contig024) [29]. The sequence from EzBioCloud for this same strain (accession no. AF490539.1) contains two errors, see text.

These data indicated that PAM 2766^T^ was genomically sufficiently distinct from all the species within the rhodococcal monophyletic sublineage containing *R. equi* to warrant separate species status.

### 16S rDNA SEQUENCE

The 16S rRNA gene sequence of PAM 2766^T^ (from both the draft genome assembly and a 16S rDNA PCR amplicon obtained using oligonucleotides primers 27F 5’-AGAGAGTTTGATCCTGGCTGGCTCAG-3’ and 1492R 5’-CGGCTACCTACCTTGTTACGACTT-3’ [28] and high-fidelity DNA polymerase Q5 [Q5® High-Fidelity DNA Polymerase, New England Biolabs]) was found to be 100% identical to that of *R. equi*. This was determined using a high-quality draft genome assembly of *R. equi* DSM 20307^T^ from our laboratory (GenBank accession no. LWTX01000024 [LWTX01000000 – contig024] [29]) (**Table S1**) as well as two other *R. equi* 16S rDNA sequences available in the databases (GenBank accession nos. X80614 for DSM 20307^T^ deposited in 1995 [30] and FJ468344 for ATCC 6939^T^ deposited in 2008). Additionally, only three of the six other species encompassed in the same rhodococcal sublineage as *R. equi* (the most distantly related on the basis of ANI/AAI, i.e. *R. subtropicus*, *R. spongiicola* and *R. xishaensis*, had 16S rRNA similarity scores below the 98.7% species demarcation standard [22, 25, 26, 31] (**Table 1**). It is not unusual among closely related rhodococci to share virtually identical 16S rRNA gene sequences [32–35], highlighting the limitations of the16S rDNA sequences for accurate differentiation of prokaryotic species [25, 36, 37].

Of note, the 16S rDNA sequence similarity dropped to 99.86% when using GenBank accession no. AF490539, designated as the reference sequence for *R. equi* DSM 20307^T^ by major databases (EzBioCloud 16S database [https://www.ezbiocloud.net/db] [38], LPSN [https://lpsn.dsmz.de/species/rhodococcus-equi] [3]). Since the 16S rDNA sequences from >100 *R. equi* WGSs (including a reference diversity set of 27 previously characterized isolates from different sources [29]; NCBI BioProject PRJNA316970]) showed all 100% similarity to LWTX01000024 (as well as X80614 and FJ468344), a caveat must be sounded as to the inaccuracy of the AF490539 sequence. AF490539 appears to contain two sequencing errors: a missing cytosine (of a series of four cytosines) between C50 and T51, and a missing guanine (of a series of five guanines) between C53 and G54.

### PHYLOGENOMIC ANALYSIS

To determine the phylogenetic position and relationships of PAM 2766^T^, we generated a core-genome phylogeny using all available genome assemblies for the type strains of each species of the genus *Rhodococcus* (as emended by Val-Calvo and Vázquez-Boland 2023 [11, 39]) with status name “correct” in the LPSN repository (https://lpsn.dsmz.de/genus/rhodococcus, accessed January 2024) (**Table S1**). The *Rhodococcoides* gen. nov. [11] circumscription, which forms a distinct, early diverging branch at the base of the rhodococcal radiation [11, 29], was excluded from the phylogeny except for the type species, *Rhodococcoides* (formerly *Rhodococcus*) f*ascians* NBRC 12155^T^, which was used as an outgroup. Using a closely related outgroup maximizes the number of core genomic markers and thus the resolution of the phylogenetic reconstruction. Core-genome orthologous genes were identified using the Get-Homologues package v. 22082022 [40] (COG and OrthoMCL clustering intersect with settings: minimal coverage 75%, maximum E-value 1e-05) and filtered to exclude recombinant alignments or alignments producing anomalous or poorly supported trees using the Get-Phylomarkers v2.4.6 tool [41]. Concatenated alignments based on the nucleotide sequence of conserved CDS (409 gene markers) and the amino acids sequence of conserved gene products (323 protein markers) were generated (**Fig. 1**). Maximum Likelihood (ML) phylogenetic trees were built using IQtree2 software version 2.2.0 [42] with the substitution models GTR+F+ASC+R5 and LG+F+R3A for the nucleotide and protein sequences, respectively.

The resulting nucleotide- and protein sequence-based *Rhodococcus* genus ML trees were strongly supported and fully consistent with each other (**Fig. 1**). Both phylogenies located PAM 2766^T^ in rhodococcal sublineage no. 2 [11] as the most closely related species to *R. equi*. The evolutionary distance between PAM 2766^T^ and *R. equi* is equivalent to or greater than that between other pairs of closely related species within other *Rhodococcus* sublineages (e.g. *Rhodococcus rhodochrous* and *Rhodococcus pyridinivorans* in sublineage 1, or *Rhodococcus opacus* and *Rhodococcus wratislaviensis* in sublineage 4 [11]) (**Fig. 1**), also supporting the classification of PAM 2766^T^ as a distinct species. This is most clearly illustrated by a core-genome ML tree of sublineage 2 *Rhodococcus* spp. in which several *R. equi* isolates representative of the diversity of the species were included for reference (**Fig. 2**). Here, while the *R. equi* isolates form a punctiform radiation due to their extremely short relative genetic distances [29], PAM 2766^T^ branches out at a significant distance in a comparable fashion to the other species encompassed in this rhodococcal sublineage. The tree also shows that PAM 2766^T^ represents the most closely related taxon to *R. equi* in the *Rhodococcus* circumscription (**Fig. 2**).

**Fig. 2.**
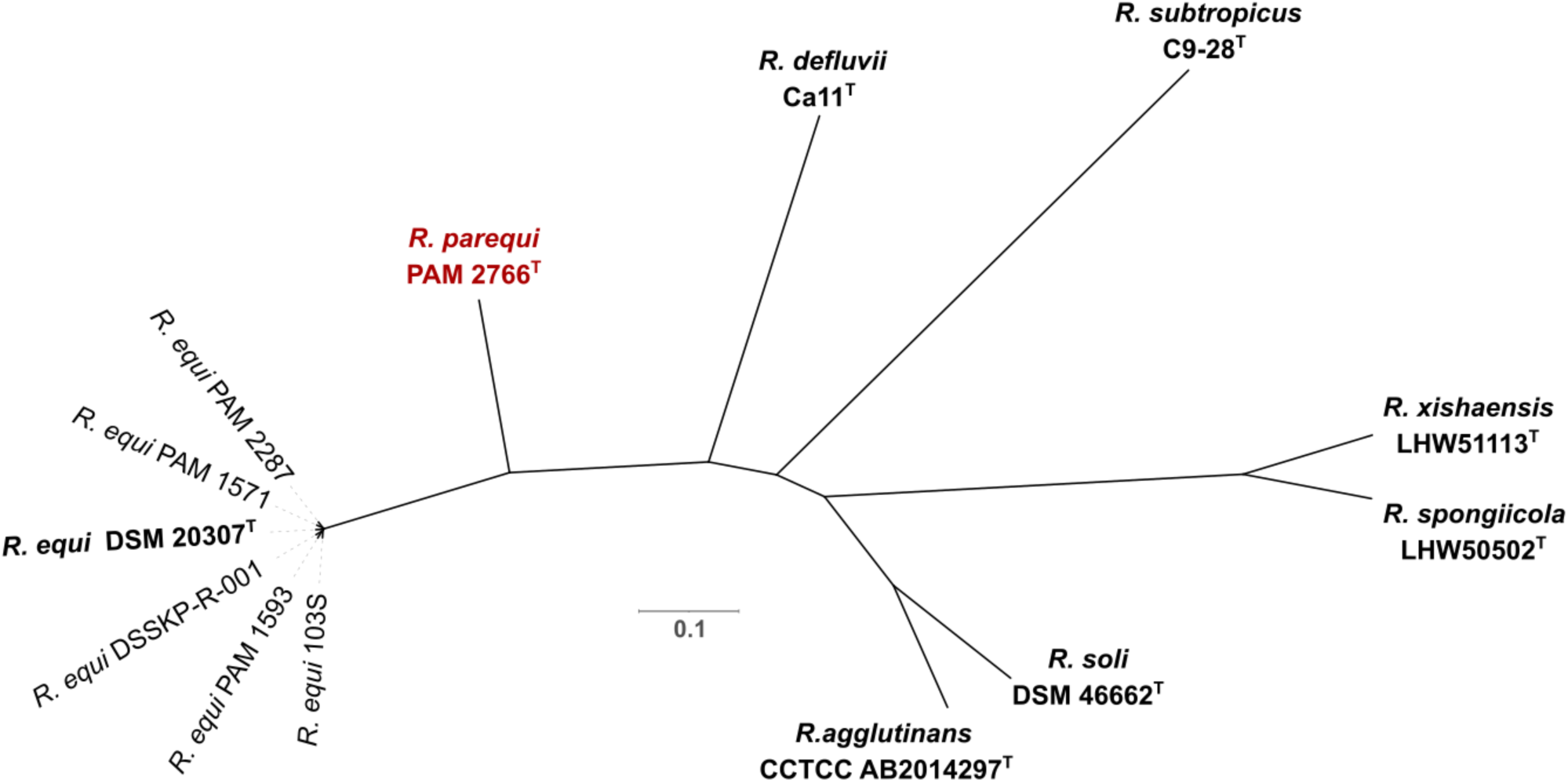
Unrooted core-genome ML tree of *Rhodococcus* monophyletic sublineage no. 2 containing *R. equi* and *R*. *parequi* PAM 2766^T^. The analysis includes a selection of *R. equi* strains representative of the genomic diversity of the species [29]. Based on a concatenated alignment of 828 core genes identified using Get-Homologues v. 22082022 [40] (orthologs identified with the COG and OrthoMCL clustering algorithms intersect using as settings minimal coverage 75%, maximum E-value 1e-05). Substitution model was GTR+F+ASC+R3. Branch support for each node is 100 (1,000 ultrafast bootstrap replicates using IQtree UFBoot).

### PHENOTYPIC CHARACTERIZATION

PAM 2766^T^ are Gram-positive, non-spore-forming coccobacilli morphologically similar to *R. equi* and display the general biochemical/physiological profile of the genus *Rhodococcus* (strictly aerobic, non-motile, catalase positive and oxidase negative). The growth characteristics are akin to those of *R. equi*, both species forming on tryptic soy agar (TSA) smooth, creamy, buff-coloured, shiny colonies that are difficult to differentiate from each other (**Fig. S1**). Grading of growth intensity/abundance in different media was TSB (VWR Chemicals 84675) > BHI (VWR Chemicals 84626) > LB (Sigma L3022) based on growth curves in an Optima BMG plate reader (**Fig. S2**). Also similar to *R. equi* [23], PAM 2766^T^ requires thiamine supplementation for growth (**Table 2**, **Fig. S3A**). Genomic analysis of *R. equi* identified the disruption of the *thiCD* locus in the thiamine biosynthesis pathway, caused by a horizontal gene acquisition event with concomitant deletion of the *thiC* gene, as the likely cause of the thiamine auxotrophy [23, 29] (**Fig. S3B**). Inspection of the genetic structure of the homologous region in PAM 2766^T^ showed it was virtually identical to that of *R. equi*, including the same *thiCD* genomic lesion (**Fig. S3B**). Since species with a complete *thiCD* locus are also present in the *Rhodococcus* radiation (e.g. *R. erythropolis* [23] [**Fig. S3B**]), the *thiCD* disruption/*thiC* deletion likely took place somewhere between the common ancestor of the genus and that of the rhodococcal sublineage encompassing *R. equi* and PAM 2766^T^. The thiamine auxotrophy of *R. equi* has been seen as an adaptive trait to the natural reservoirs of this species, manure-rich soil and the large intestine, where microbiota-derived thiamine is likely to be readily accessible [8]. We surmise a similar explanation may apply to PAM 2766^T^.

**Table 2.** Main phenotypic characteristics of *R. parequi* PAM 2766^T^ and *R. equi.* Results: positive, +; weakly positive, w; negative, −. See table footnotes for methods and **Fig. 4** for an overview of Biolog’s Phenotype MicroArray^TM^ results.

|  | *R. parequi*<br>PAM 2766<sup>T</sup> | *R. equi*<br>DSM 20307<sup>T</sup> | *R. equi* 103S<br>(PAM 1126) |
| --- | --- | --- | --- |
| Growth temperature range (°C) <sup>a</sup> | 5 <sup>b</sup> – 45 | 10 – 45 <sup>c</sup> | 10 – 45 |
| pH range <sup>c</sup> | 5 – 10 | 5 – 10 | 5 – 10 |
| NaCl tolerance range <sup>d</sup> | 1 – 3 | 1 – 4 | 1 – 4 <sup>e</sup> |
| Anaerobic growth <sup>f</sup> | – | – | – |
| Motility <sup>g</sup> | – | – | – |
| Catalase | + | + | + |
| Oxidase | – | – | – |
| Urease <sup>h</sup> - 24h | + | w | – |
| - 48h | + | + | w |
| Thiamine auxotrophy <sup>i</sup> | + | + | + |
| CAMP-like test <sup>j</sup> | + | + | + |
| API Coryne profile | 1110004 | 1110004 | 1110004 |
| Phenotype MicroArray<sup>TM</sup> |  |  |  |
| C-sources |  |  |  |
| Tween 20 | – | + | + |
| 4-hydroxybenzoic acid | – | + | + |
| b-hydroxybutyric acid | – | + | + |
| N-sources |  |  |  |
| L-cysteine | – | + | + |
| L-lysine | – | + | + |
| L-citruline | – | + | + |
| L-ornithine | – | + | + |
| Thymine | – | + | + |
| e-amino-N-caproic acid | – | + | + |
| d-amino-N-valeric acid | – | + | + |

<sup>a</sup> Tested on TSA plates.
<sup>b</sup> Growth at 5°C was observed after 10 days.
<sup>c</sup> Differs from emendation of *R. equi* (Magnusson 1923) Goodfellow and Alderson 1977 by Goodfellow *et al.* 1982 [55] where a temperature growth range of 5 to 40 °C was reported.
<sup>d</sup> Determined using growth curve assays in TSB at 30 °C using an Optima BMG plate reader (48-well plates with 400µl medium/well, 200 rpm shaking, readings every 30 min). The pH of the medium was adjusted at 1.0 intervals with phosphate (pH 2.0–7.0), Tris-HCl (pH 7.0–9.0) or sodium bicarbonate (pH 9.0–11.0). NaCl concentrations between 0 and 10 % (w/v) were tested at 1% intervals.
<sup>e</sup> Differs from the results reported by Lee *et al.* (2019) [6] using ISP 2 agar.
<sup>f</sup> Determined in TSA incubated in an anaerobic chamber with AnaeroGen<sup>TM</sup> sachet (Thermo Fisher) at 30 °C for 20 days.
<sup>g</sup> Determined in TSB 0.3% agar incubated at 30 °C for 14 days.
<sup>h</sup> Determined using Christensen’s Urea Agar as per ASM protocols (https://asm.org/getattachment/ac4fe214-106d-407c-b6c6-e3bb49ac6ffb/urease-test-protocol-3223.pdf) modified by replacing dextrose (which *R. equi* does not use, or uses inefficiently) with 25 mM sodium lactate (preferred carbon source that promotes vigorous growth of *R. equi* [29]).
<sup>i</sup> Determined *R. equi* mineral medium [56] as modified in refs. [23, 29] (mReMM) supplemented with 20 mM lactate using growth curve assays in an Optima BMG plate reader (see footnote d and **Fig. S3**).
<sup>j</sup> Synergistic hemolysis with sphingomyelinase C-producing bacteria determined in Columbia 5% sheep blood agar using *L. ivanovii* as indicator strain as previously described [13, 56].

A search for phenotypic differentiation markers found that PAM 2766^T^ can grow at 5° C, evident after 10 days of incubation, in contrast to *R. equi* DSM 20307^T^ (and the *R. equi* reference genome strain 103S [PAM 1126] as a representative of the other major phylogenomic subdivision of the species [29]) (**Table 2**). PAM 2766^T^ also gave a clearly positive urease test after 24 h at 30°C, while *R. equi* required a longer incubation (minimum of 48 h) and the reaction was generally weaker (**Table 2**). It is worth noting that the urease test was in all cases negative using the API Coryne strips (Biomerieux). The reason for the discrepancy is that in the API Coryne gallery urease activity is tested under anaerobic conditions, preventing the growth of the strictly aerobic *Rhodococcus* bacteria.

Phenotype MicroArray^TM^ (Biolog Inc.) screens for carbon (PM1 and PM2A plates) and nitrogen (PM3B plates) sources were used to identify additional differential phenotypic markers. PAM 2766^T^ utilized, with different efficiency, 26 out of the 190 tested carbon sources. Like *R. equi* [23], PAM 2766^T^ seems to assimilate carbon mainly through lipid metabolism, growing vigorously on different organic acids as a sole carbon source, including lactate, acetate and sorbate (**Fig. 3, Table S2**). PAM 2766^T^ could be differentiated from *R. equi* by the inability to growth on Tween 20 (polyoxyethylene sorbitan monolaurate), 4-hydroxybenzoic acid and β-hydroxybutyric acid. Tween 20 has been reported to be toxic for some *Rhodococcus* spp. while it is utilized by other rhodococci such as *Rhodococus jostii* or *Rhodococcus opacus* [43, 44] in addition to *R. equi* [23]. PAM 2766^T^ assimilated a wide range of nitrogen sources, 41 out of 95 compounds tested in the PM3B plates (**Fig. 3, Table S2**). As main differences with *R. equi*, PAM 2766^T^ was unable to grow on L-cysteine, L-lysine, L-citrulline, L-ornithine, cytosine, ε-amino-N-caproic acid and δ-amino-N-valeric acid. The relevant phenotypic characteristics of PAM 2766^T^ are summarized in **Table 2** and below in the species description.

**Fig. 3.**
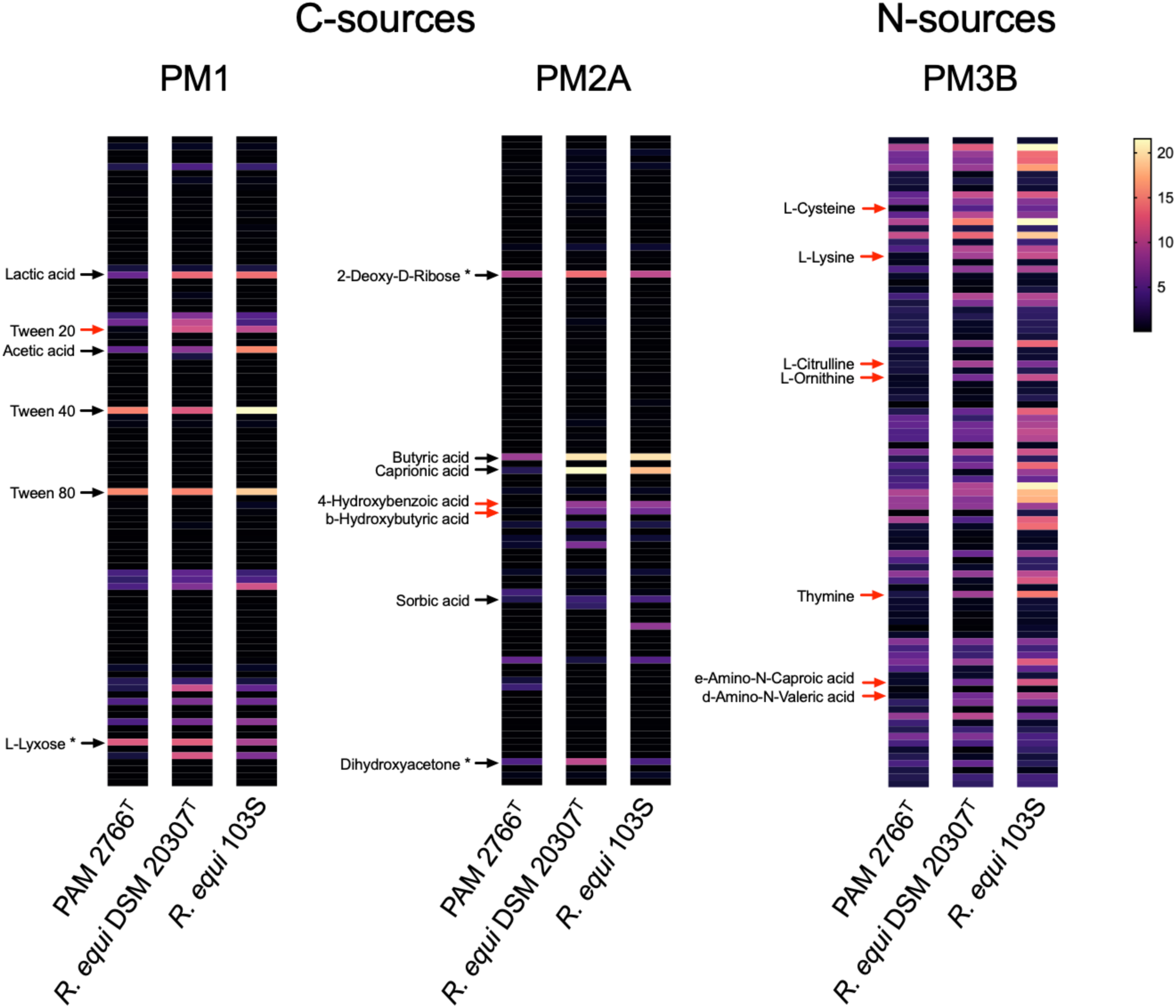
Heat map of Phenotype MicroArray^TM^ (PM) results for carbon and nitrogen source utilization by PAM 2766^T^, *R. equi* DSM 20307^T^ and *R. equi* 103S (PAM 1126). Bacterial inocula were grown at 30°C in TSB until the stationary phase and then suspended in mReMM mineral medium and transferred to the PM plates. Incubabion was performed at 30°C with OD_590_ monitored every 15 min for 48 hours in an OmniLog reader. Strains were tested in duplicate and results were analysed using OmniLog software. Maximum growth is represented in graded colours from lowest (black) to highest (yellow). Red arrows indicate differential utilization of a substrate between PAM 2766^T^ and *R. equi.* Black arrows in the PM1 and PM2A plates indicate a carbon source utilized by the three tested bacteria (see **Table S2** for detailed results). Asterisks indicate false positive reactions in the PM2A plate previously reported in ref. [23].

### VIRULENCE

Although PAM 2766^T^ was isolated from soil and we confirmed its genome lacked any of the plasmid virulence determinants of *R. equi* [45], this did not exclude it might have pathogenic potential. To explore this, we tested PAM 2766^T^ in macrophage infection essays. The ability to survive and multiply within macrophages is at the basis of the infectivity of *R. equi* and other related pathogenic actinomycetes [46, 47]. Infection assays were performed in J774A.1 mouse macrophages cultured until confluence using a vancomycin protection essay as previously described [18, 23] (see **Fig. 4** legend for experimental details). Virulent *R. equi* 103S [23] and the non-virulent isogenic derivative 103S^−^ (obtained by curation of the pVAPA virulence plasmid [48] required for intramacrophage survival [18, 49, 50]) were used as positive and negative controls. *R. equi* 103S showed the expected behaviour, with significant intramacrophage proliferation during the infection time course. In contrast, PAM 2766^T^ intracellular numbers progressively declined, mirroring the behaviour of the pVAPA-cured, non-virulent *R. equi* 103S^−^ strain (**Fig. 4**). These data support the notion that PAM 2766^T^ is not primarily pathogenic and most likely represents an environmental saprotrophic species.

**Fig. 4.**
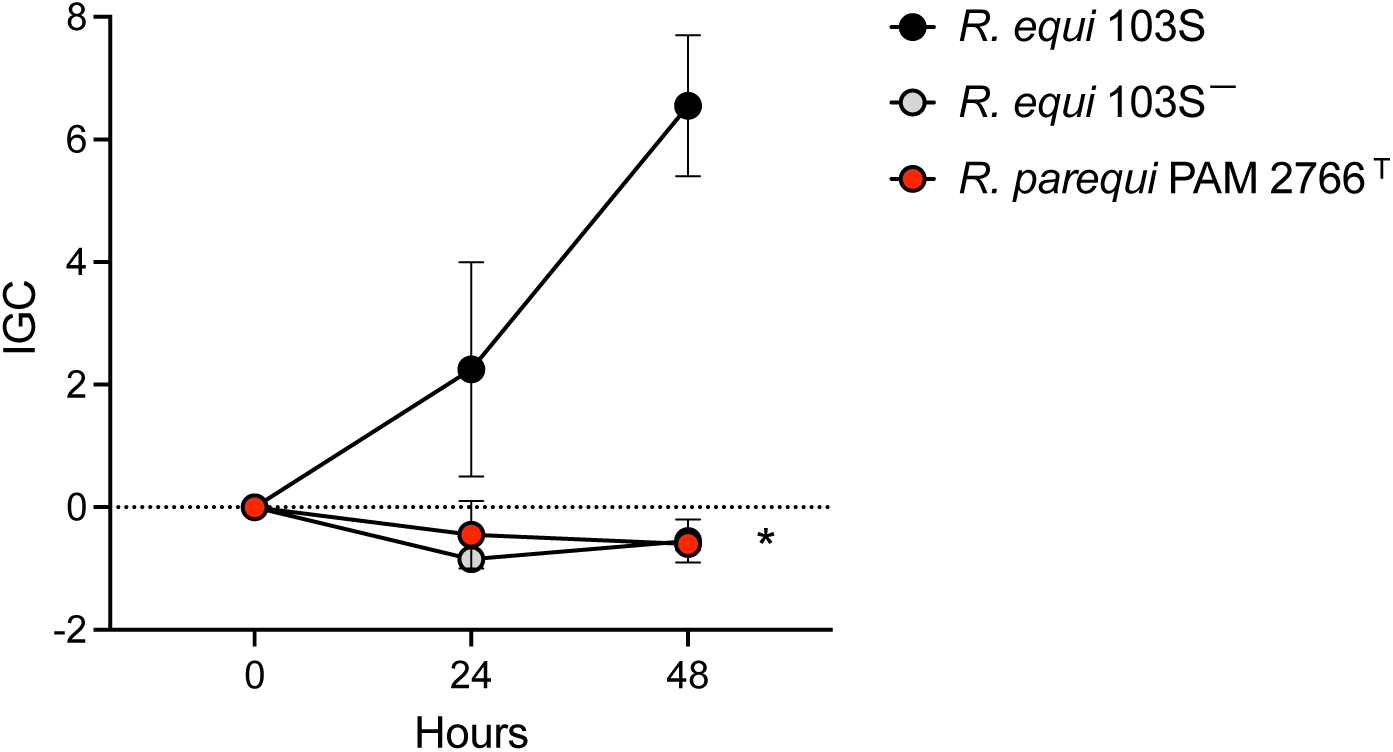
Macrophage infections. Intracellular proliferation phenotype of *R. parequi* PAM 2766^T^, virulent *R. equi* 103S (harbouring the pVAPA virulence plasmid that promotes intramacrophage replication [18, 53, 54]) and non-virulent *R. equi* 103S^−^ (isogenic pVAPA-cured derivative unable to proliferate intracellularly [52]) in the J774A.1 murine macrophage-like cell line. J774A.1 cells (ATCC, <10 passages) were cultured in 24-well plates at 37 °C with 5% CO_2_ in RPMI supplemented with pyruvate and glutamine and 10 % foetal bovine serum (Lonza) until confluence (≈2×10^5^ cells/well). Macrophage monolayers were infected at 10∶1 multiplicity with washed bacteria from an exponential culture at 37°C in TSB (OD_600_ ≈1.0). They were then centrifuged for 3 min at 172×g, incubated for 60 min at 37°C, washed three times with PBS to remove nonadherent bacteria, and incubated in RPMI supplemented with 5µg/µl vancomycin to prevent extracellular bacterial growth. After 1 h of incubation with vancomycin (*t* = 0) and at 24 and 48 h thereafter, macrophage monolayers were washed with PBS, lysed with 0.1% triton X-100 for 3 min, and intracellular bacterial counts determined by plating appropriate dilutions of the cell lysates onto TSA. As the intracellular bacterial population at a given time point depends on initial numbers, for data comparability between strains, bacterial intracellular kinetics data are expressed as a normalized Intracellular Growth Coefficient (IGC) = (IB*_t_* _= *n*_−IB*_t_* _= 0_)/IB*_t_* _= 0_, where IB*_t_* _= *n*_ and IB*_t_* _= 0_ are the intracellular bacterial numbers at a specific time point *t* = *n* and *t* = 0, respectively. Positive IGC indicates proliferation, negative values reflect a decrease in the intracellular bacterial population. Bacterial counts per well at *t* = 0: 103S^+^, 2.55±0.91×10^3^; 103S^−^, 1.75±0.17×10^3^; *R. parequi* PAM 2766^T^, 1.67±0.18×10^3^. Means of two independent duplicate experiments ±SEM. Asterisks denote significant differences at *t* = 48 h with *R. equi* 103S ( *P*≤0.001, two-way ANOVA).

### TAXONOMIC CONCLUSION

The clear cut phylogenetic/genomic separation from the rest of the rhodococci and the identification of phenotypic markers allowing its differentiation from the closely related *R. equi* in our opinion justify the consideration of strain PAM 2766^T^ as a distinct *Rhodococcus* species. Owing to the remarkable morphological and biochemical similarities with *R. equi*, close phylogenetic relationship and similar source ecosystem, we propose for this novel species the name *Rhodococcus paraequi*. However, according to appendix 9A(2) of the International Code of Nomenclature of Prokaryotes (ICNP) [57] the final vowel of the prefix *para* has to be elided, giving *Rhodococcus parequi* sp. nov.

### DESCRIPTION OF *RHODOCOCCUS PAREQUI* SP. NOV

Etymology: *Rhodococcus parequi* (par.e’qui. Gr. prep. *para*, beside, alongside of, near, like; L. gen. masc. n. *equi*, of a horse, specific epithet; N.L. gen. n. *parequi*, resembling *Rhodococcus equi*).

Members of this species are Gram-positive, strictly aerobic, non-spore-forming, non-motile, catalase positive, oxidase negative coccobacilli. *R. parequi* grows well in standard solid culture media such as TSA, BHI agar, or LB agar. After 24 to 48 h incubation at 30° C, it forms smooth, shiny, creamy to buff-coloured colonies that tend to coalesce over time. Colonies are morphologically very similar to those of *R. equi*. No haemolysis is observed on Columbia 5% sheep blood agar but gives a positive synergistic lytic (CAMP-like) reaction with sphingomyelinase C-producing bacteria such as *L. ivanovii* or *S. aureus* [13, 52]. This ability is shared with *R. equi* and is the phenotypic expression of cholesterol oxidase production [12]. *R. parequi* also tests positive to a PCR targeting the *R. equi* cholesterol oxidase gene *choE* [14], considered up to know to be a species-specific identification marker for *R. equi* [14–17]. Optimal growth temperature is between 30 and 37 °C. Tolerates NaCl concentrations up to 3 % (w/v) and pH values from 5 to 9, with 7 as pH optimum. Tests positive for urea hydrolysis, nitrate reduction, acetoin production, tryptophane deaminase, α-glucosidase, and alkaline phosphatase. It can be distinguished from *R. equi* (as tested with DSM 20307^T^ and 103S [PAM 1126]) by a rapid urease positive reaction (at 24 h incubation), ability to grow at 5 °C, and inability to utilize Tween 20, 4-hydroxybenzoate and β-hydroxybutare as a carbon source, and L-citrulline, L-ornithine, cytosine, ε-amino-N-caproic acid and δ-amino-N-valeric acid as a nitrogen source. The draft genome of strain PAM 2766^T^ is 5.3 Mbps in size with a digital G+C content of 68.98 mol %. Phylogenomically *R. parequi* belongs to sublineage no. 2 of the *Rhodococcus* radiation (as defined in ref. [11]) together with *R. agglutinans*, *R. defluvii*, *R. equi, R. soli* and *R. subtropicus*. Genomic similarity indices with its most closely related species *R. equi* are ANI = 88.60 and AAI = 92.35. *R. parequi* is avirulent in macrophage infection assays and is assumed to be a non-pathogenic environmental saprophyte.

The type strain is PAM 2766^T^ (CETC 30995^T^ = NCTC 14987^T^) isolated from equine farm soil in Normandy, France.

## Acknowledgements

We would like to thank Enrico Tatti and Barry Bogner from Biolog Inc. for their help and for facilitating the collaboration with C.V. and F.D. D. Lewis is gratefully acknowledged for her initial contributions in setting up our global *R. equi* laboratory collection. Thanks are also due to M. Göker for advising that *R. paraequi* should be named *R. parequi* as per ICNP rules. This study was supported by the Horserace Betting Levy Board (HBLB projects nos. vet/prj/796 and vet/prj/814).

## Authors contributions

J.V-B., conceptualization, funding acquisition, formal analysis, and writing up – original draft, review and editing; J.V-C., investigation, formal analysis, visualization and writing up – original draft, review and editing; C.V and F.D., investigation, resources; F.D. and S.P, investigation, resources; M.S., investigation, formal analysis, visualization and writing up – original draft, review and editing. All authors read, commented on and approved the manuscript.

## Abbreviations

AAI: average amino acids identity
ANI: average nucleotide identity
BHI: brain-heart infusion
BHIA: BHI agar
bp: base pairs
CAMP: acronym of Christie-Atkins-Munch-Petersen test of synergistic haemolysis on sheep blood agar with sphingomyelinase C-producing *S. aureus*
CDS: coding sequence
CECT: Colección Española de Cultivos Tipo - Spanish Type Culture Collection
dDDH: digital DNA-DNA hybridization
GTR: General Time-Reversible with empirical base frequencies
ICNP: International Code of Nomenclature of Prokaryotes
LB: Luria-Bertani
LPSN: Lists of Prokaryotic names with Standing in Nomenclature (https://www.bacterio.net)
LG: Le-Gascuel general amino acid replacement matrix
Mbp: mega base pairs
ML: maximum likelihood
NCTC: National Collection of Type Cultures
OGRI: overall genomic relatedness indices
PAM: “Patogénesis Microbiana” isolate collection of Vazquez-Boland’s laboratory
ReMM: *R.equi* mineral medium
SEM: standard error of the mean
TSA: tryptic soy agar
TSB: tryptic soy broth
PCR: polymerase chain reaction
WGS: whole genome sequencing
+ASC: ascertainment bias correction
+F: empirical base frequencies
+R: FreeRate model categories.

