## Supplemental figures and tables for "*Rhodococcus parequi* sp. nov., a new species isolated from equine farm soil closely related to the pathogen *Rhodococcus equi*"

### SUPPLEMENTAL MATERIAL

#### Supplemental Figures

48h 30ºC | + 3d RT

*R. parequi* PAM 2766$^T$

*R. equi* DSM 20307$^T$

*R. equi* 103S

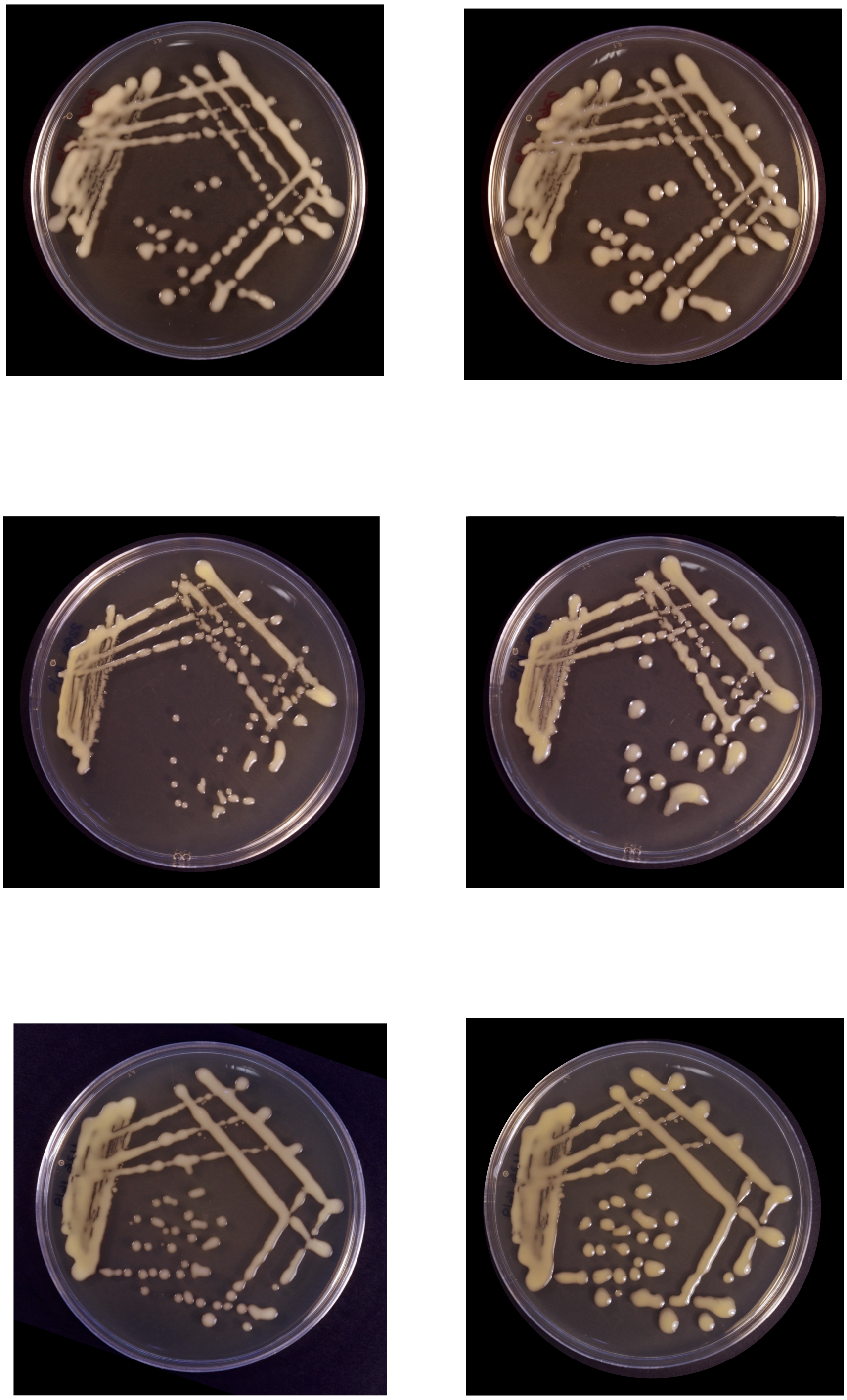

**Fig. S1.** Colony morphology of *R. parequi* PAM 2766 $^T$ compared to *R. equi* (type strain DSM 20307$^T$ and reference genome strain 103S (PAM 1126), each belonging to one of the two main phylogenomic subdivisions of the species [29]. TSA plates were grown at 30 ºC for 48 h (left) and then left three additional days at room temperature.

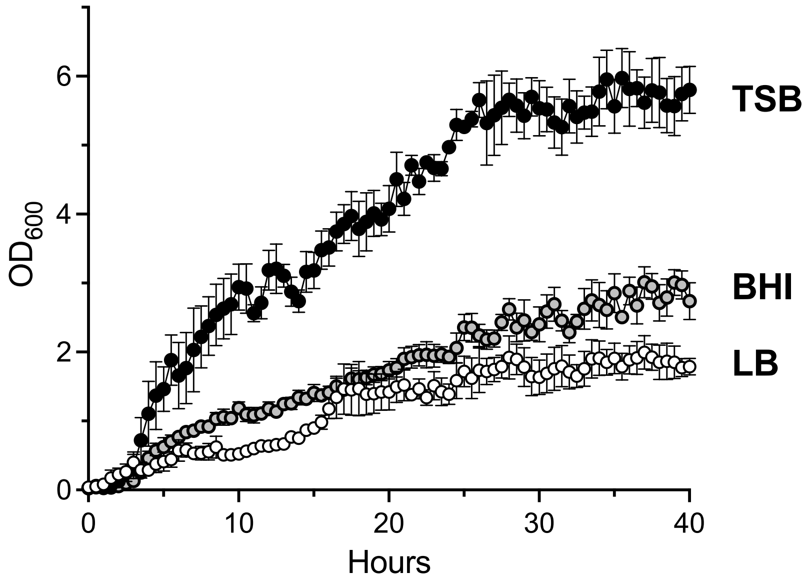


**Fig. S2**. Growth dynamics of PAM 2766$^T$ in different culture media: tryptic soy broth (TSB), brain-heart infusion (BHI), Luria-Bertani broth (LB). Growth was monitored in 48-well plates (Costar) by measuring the $OD_{600}$ every 30 min in an automated plate reader (Optima apparatus, BMG Labtech) during incubation at 30 ºC with 400 rpm shaking. Bacterial cells were obtained from an overnight culture in BHI incubated at 30 ºC, washed in PBS and resuspended in the appropriate medium to an optical density at 600nm ($OD_{600}$) ≈ 0.05. Wells were inoculated in duplicate using 400µl aliquots.

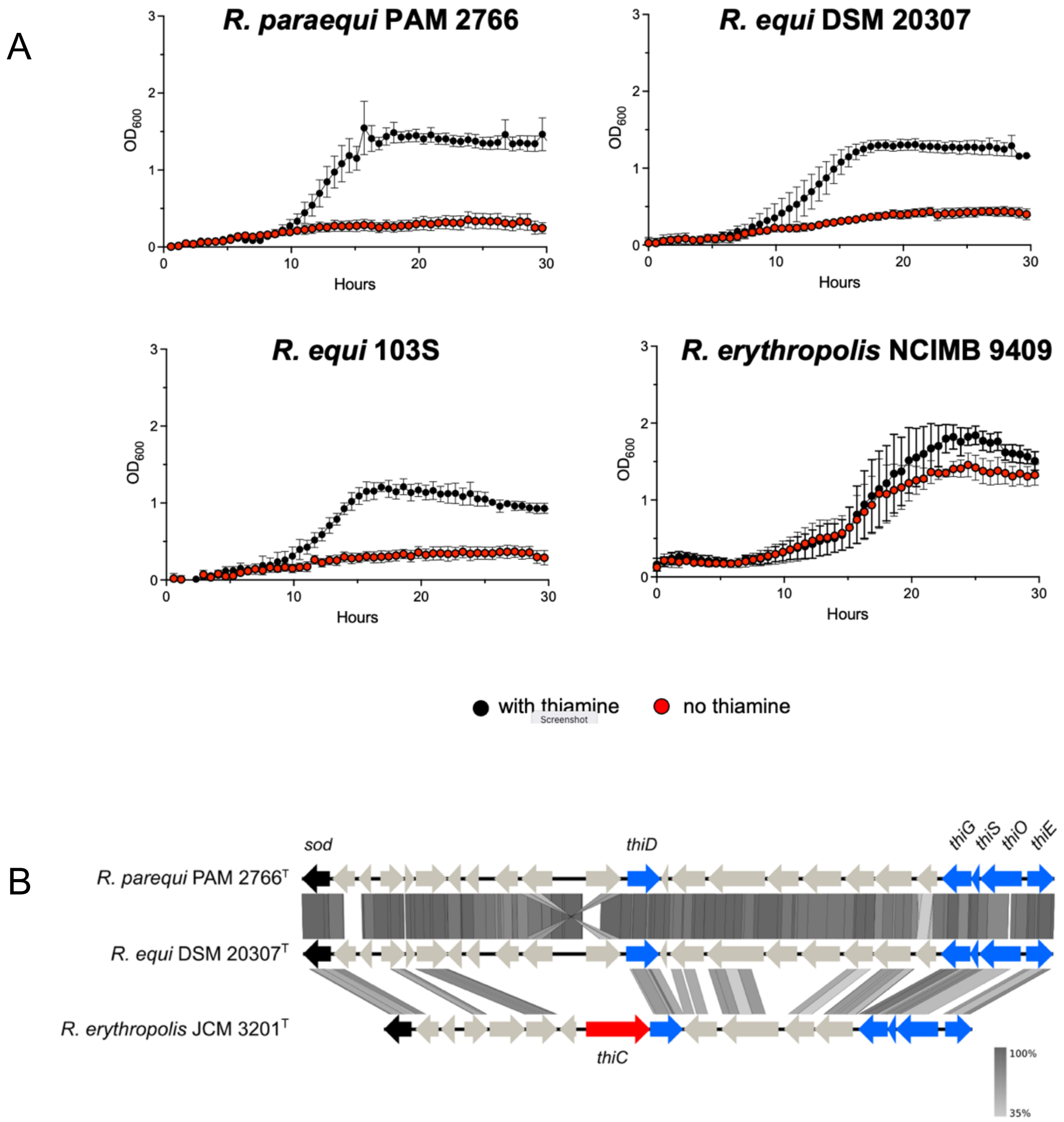


**Fig. S3.** Thiamine dependency of *R. parequi* PAM 2766$^{T}$. (A) Growth curves of *R. parequi* PAM 2766$^{T}$, *R. equi* (two strains strains representing the two main lineages of the species, DSM 20307$^{T}$ and 103S) and a representative strain of *R. erythropolis* in 20 mM lactate mReMM mineral medium [29] with or without 0.1 mM thiamine supplementation. Bacteria were grown overnight in TSB, collected by centrifugation, washed in PBS and resuspended in mReMM to $OD_{600} \approx 0.05$. Duplicate 400µl aliquots of each bacterial suspension were distributed in a 48-well plate (Costar). Plates were incubated in an automated plate reader (Omega, BMG Labtech) at 30ºC with double orbital shaking (400 rpm) and growth was monitored every 30 min. Mean of at least three duplicate experiments ± SEM. Chemicals were purchased from Sigma. (B) TBlastX alignment of the chromosomal *thiCD-GSOE* thiamine biosynthesis locus of *R. equi* DSM 20307$^{T}$ compared to that of *R. parequi* PAM 2766$^{T}$ and *R. erythropolis* JCM 3201$^{T}$. Note that the genomic region is virtually identical in *R. equi* and *R. parequi*. Both species lack the *thiC* gene and are thiamine auxotrophs in contrast to *R. erythropolis* JCM 3201$^{T}$ (as an example of a *Rhodococcus* sp. not requiring thiamine for efficient growth).

**Table S1.** Genome assemblies of type strains used in this study.

| Species | Strain | NCBI assembly information | | | | | |
|---|---|---|---|---|---|---|---|
| | | G+C content (%) | Size (Mbps) | No. of contigs/scaffolds | Completeness (%) | Contamination (%) | Accession number |
| *Rhodococcoides fascians* | NBRC 12155 | 64.5 | 5.8 | 36 | 99.5 | 1.46 | GCF_001894785.1 |
| *Rhodococcus aetherivorans* | DSM 44752 | 70.5 | 6.4 | 216 | 79.86 | 0.99 | GCF_011058165.1 |
| *Rhodococcus agglutinans* | CFHS 0262 | 69.0 | 5.4 | 22 | 99.5 | 1.12 | GCF_004011865.1 |
| *Rhodococcus antarcticus* | 75 | 73.0 | 3.9 | 4 | 84.36 | 3.27 | GCF_026153295.1 |
| *Rhodococcus coprophilus* | NCTC 10994 | 67.0 | 4.6 | 1 | 96.8 | 1.04 | GCF_900478115.1 |
| *Rhodococcus defluvii* | Ca11 | 68.5 | 5.1 | 267 | 98.24 | 0.87 | GCF_000738775.1 |
| *Rhodococcus electrodiphilus* | LMG 29881 | 70.5 | 5.5 | 225 | 98.91 | 0 | GCF_030509825.1 |
| *Rhodococcus equi* | DMS 20307 | 69.0 | 5.2 | 37 | 98.33 | 0.99 | GCF_002094305.1 |
| *Rhodococcus erythropolis* | JCM 3201 | 62.5 | 6.7 | 3 | 94.82 | 4.28 | GCF_003990875.1 |
| *Rhodococcus globerulus* | NBRC 14531 | 61.5 | 6.7 | 30 | 99.19 | 0.78 | GCF_001894805.1 |
| *Rhodococcus gordoniae* | NCTC 13296 | 68.0 | 4.9 | 3 | 98.86 | 0.33 | GCF_900455725.1 |
| *Rhodococcus indonesiensis* | CSLK01-03 | 70.0 | 5.5 | 282 | 98.58 | 0 | GCF_030360185.1 |
| *Rhodococcus jostii* | DSM 44719 | 67.0 | 9.9 | 6 | 97.82 | 2.46 | GCF_900105375.1 |
| *Rhodococcus koreensis* | DSM 44498 | 67.5 | 10.3 | 9 | 99.35 | 4.95 | GCF_900105905.1 |
| *Rhodococcus maanshanensis* | DSM 44675 | 69.0 | 5.7 | 61 | 97.89 | 2.62 | GCF_900109405.1 |
| *Rhodococcus marinonascens* | NBRC 14363 | 64.5 | 4.9 | 156 | 95.52 | 1.1 | GCF_001894885.1 |
| *Rhodococcus opacus* | DSM 43205 | 67.0 | 9.0 | 38 | 99.5 | 3.59 | GCF_910591545.1 |
| *Rhodococcus oryzae* | NEAU-CX67 | 69.0 | 5.4 | 31 | 98.22 | 2.29 | GCF_005049235.1 |
| *Rhodococcus oxybenzonivorans* | S2-17 | 65.5 | 8.0 | 4 | 98.85 | 1.74 | GCF_003130705.1 |
| *Rhodococcus phenolicus* | DSM 44812 | 68.5 | 6.3 | 232 | 97.45 | 2.16 | GCF_001646785.1 |
| *Rhodococcus pseudokoreensis* | R79 | 67.5 | 9.9 | 6 | 99.35 | 3.24 | GCF_017068395.1 |
| *Rhodococcus pyridinivorans* | DSM 44555 | 68.0 | 5.3 | 3 | 96.81 | 2.53 | GCF_900105195.1 |
| *Rhodococcus rhodnii* | ATCC 35071 | 69.5 | 4.5 | 4 | 92.35 | 1.03 | GCF_008011915.1 |
| *Rhodococcus rhodochrous* | NCTC 10210 | 68.0 | 5.3 | 1 | 99.44 | 0.99 | GCF_900187265.1 |
| *Rhodococcus ruber* | NBRC 15591 | 70.5 | 5.3 | 56 | 99.27 | 0.37 | GCF_001894945.1 |
| *Rhodococcus soli* | DSM 46662 [a] | 68.8 | 5.4 | 20 | 98.71 | 0.68 | JBDLNU000000000 |
| *Rhodococcus spelaei* | C9-5 | 69.0 | 4.8 | 18 | 97.11 | 1.21 | GCF_006704125.1 |
| *Rhodococcus spongiicola* | LHW 50502 | 66.5 | 4.0 | 29 | 96.65 | 1.3 | GCF_004011835.1 |

[a] The genome sequence of *R. soli* DSM 46662$^{T}$ was unavailable at the time of the study and was determined *de novo*.

**Table S1.** (cont.)

| Species | Strain | NCBI assembly information | | | | | |
|---|---|---|---|---|---|---|---|
| | | G+C content (%) | Size (Mbps) | number of contigs/scaffolds | Completeness (%) | Contamination (%) | Accession number |
| *Rhodococcus subtropicus* | C9-28 | 69 | 4.4 | 67 | 98.2 | 0.65 | GCF_005434945.1 |
| *Rhodococcus triatomae* | DSM 44892 | 68.5 | 4.8 | 1 | 97.77 | 1.21 | GCF_014217785.1 |
| *Rhodococcus tukisamuensis* | JCM 11308 | 70.0 | 5.5 | 40 | 97.62 | 1.66 | GCF_900101735.1 |
| *Rhodococcus wratislaviensis* | NCTC 13229 | 67.5 | 7.8 | 29 | 99.19 | 1.24 | GCF_900455735.1 |
| *Rhodococcus xishaensis* | LHW51113 | 66.5 | 3.7 | 22 | 96.71 | 0.8 | GCF_004011825.1 |
| *Rhodococcus yananensis* | FBM 22-1 | 68.5 | 4.2 | 178 | 94.65 | 0.91 | GCF_020515525.1 |
| *Rhodococcus zopfii* | NBRC 100606 | 68 | 6.3 | 146 | 98.61 | 3.24 | GCF_001895025.1 |

**Table S2.** Phenotype MicroArray data.

| Plate | Well | Chemical | Mode of action | Maximmun growth rate | | | | | |
|---|---|---|---|---|---|---|---|---|---|
| | | | | PAM 2766 | | *R. equi* DSM 20307$^T$ | | *R. equi* 103S PAM 1126 | |
| | | | | Mean | SEM | Mean | SEM | Mean | SEM |
| PM1 | A01 | Negative Control | C-Source, negative contro | 0.1650 | 0.0550 | 0.2350 | 0.0550 | 0.1700 | 0.0100 |
| PM1 | A02 | L-Arabinose | C-Source, carbohydrate | 1.3200 | 0.3800 | 1.3400 | 0.2400 | 0.9600 | 0.2600 |
| PM1 | A03 | N-Acetyl-D-Glucosamine | C-Source, carbohydrate | 0.2000 | 0.0800 | 0.2250 | 0.0050 | 0.3300 | 0.1600 |
| PM1 | A04 | D-Saccharic acid | C-Source, carboxylic acid | 0.1100 | 0.0700 | 0.1950 | 0.0650 | 0.2050 | 0.0250 |
| PM1 | A05 | Succinic acid | C-Source, carboxylic acid | 3.3000 | 1.1900 | 6.0750 | 0.3750 | 4.8950 | 4.7050 |
| PM1 | A06 | D-Galactose | C-Source, carbohydrate | 0.1350 | 0.0050 | 0.2350 | 0.0250 | 0.1950 | 0.0350 |
| PM1 | A07 | L-Aspartic acid | C-Source, amino acid | 0.3050 | 0.2750 | 1.1650 | 1.0650 | 0.6750 | 0.5850 |
| PM1 | A08 | L-Proline | C-Source, amino acid | 0.2050 | 0.1550 | 0.4450 | 0.3350 | 0.5250 | 0.4550 |
| PM1 | A09 | D-Alanine | C-Source, amino acid | 0.3350 | 0.3150 | 0.2900 | 0.1600 | 0.1300 | 0.0900 |
| PM1 | A10 | D-Trehalose | C-Source, carbohydrate | 0.1400 | 0.0400 | 0.4150 | 0.1250 | 0.1250 | 0.0050 |
| PM1 | A11 | D-Mannose | C-Source, carbohydrate | 0.1250 | 0.0650 | 0.3300 | 0.0000 | 0.1300 | 0.0300 |
| PM1 | A12 | Dulcitol | C-Source, carbohydrate | 0.0550 | 0.0650 | 0.1350 | 0.0150 | 0.5000 | 0.4100 |
| PM1 | B01 | D-Serine | C-Source, amino acid | 0.2200 | 0.0400 | 0.1750 | 0.0250 | 0.1800 | 0.0200 |
| PM1 | B02 | D-Sorbitol | C-Source, carbohydrate | 0.1200 | 0.0500 | 0.2050 | 0.0450 | 0.5800 | 0.3800 |
| PM1 | B03 | Glycerol | C-Source, carbohydrate | 0.0550 | 0.0150 | 0.2700 | 0.0300 | 0.3500 | 0.0200 |
| PM1 | B04 | L-Fucose | C-Source, carbohydrate | 0.1550 | 0.0050 | 0.4000 | 0.1300 | 0.2500 | 0.0600 |
| PM1 | B05 | D-Glucuronic acid | C-Source, carboxylic acid | 0.2150 | 0.1550 | 0.2950 | 0.1450 | 0.1750 | 0.0450 |
| PM1 | B06 | D-Gluconic acid | C-Source, carboxylic acid | 0.1500 | 0.0900 | 0.2550 | 0.0450 | 0.1950 | 0.0750 |
| PM1 | B07 | D,L-a-Glycerol Phosphate | C-Source, carbohydrate | 0.1000 | 0.0300 | 0.1900 | 0.0300 | 0.1600 | 0.0500 |
| PM1 | B08 | D-Xylose | C-Source, carbohydrate | 2.5450 | 0.8550 | 2.1900 | 0.6700 | 2.7350 | 0.0450 |
| PM1 | B09 | L-Lactic acid | C-Source, carboxylic acid | 8.0100 | 0.6900 | 19.2250 | 2.0350 | 19.7850 | 0.3450 |
| PM1 | B10 | Formic acid | C-Source, carboxylic acid | 0.0900 | 0.0600 | 0.1750 | 0.0650 | 0.0850 | 0.0650 |
| PM1 | B11 | D-Mannitol | C-Source, carbohydrate | 0.1000 | 0.0600 | 0.1700 | 0.0500 | 0.1250 | 0.0650 |
| PM1 | B12 | L-Glutamic acid | C-Source, amino acid | 0.0150 | 0.0150 | 0.9200 | 0.2400 | 0.1550 | 0.0150 |
| PM1 | C01 | D-Glucose-6-Phosphate | C-Source, carbohydrate | 0.1400 | 0.0600 | 0.2150 | 0.0750 | 0.1600 | 0.0400 |
| PM1 | C02 | D-Galactonic acid-g-Lactone | C-Source, carboxylic acid | 0.0950 | 0.0150 | 0.1000 | 0.0100 | 0.0600 | 0.0000 |
| PM1 | C03 | D,L-Malic acid | C-Source, carboxylic acid | 5.1850 | 1.6850 | 9.7950 | 0.4550 | 6.7350 | 0.6450 |
| PM1 | C04 | D-Ribose | C-Source, carbohydrate | 8.5050 | 2.9750 | 14.7300 | 4.2700 | 5.7600 | 0.8700 |
| PM1 | C05 | Tween 20 | C-Source, fatty acid | 0.7400 | 0.2100 | 16.4200 | 0.3000 | 14.6500 | 1.3500 |
| PM1 | C06 | L-Rhamnose | C-Source, carbohydrate | 0.4850 | 0.3250 | 0.4350 | 0.1850 | 0.2300 | 0.0200 |
| PM1 | C07 | D-Fructose | C-Source, carbohydrate | 0.1300 | 0.0100 | 0.3450 | 0.1550 | 0.3400 | 0.1700 |
| PM1 | C08 | Acetic acid | C-Source, carboxylic acid | 7.9200 | 0.0000 | 10.5500 | 0.1800 | 21.7300 | 0.2100 |
| PM1 | C09 | a-D-Glucose | C-Source, carbohydrate | 0.0800 | 0.0300 | 2.7850 | 0.3750 | 0.1700 | 0.0400 |
| PM1 | C10 | Maltose | C-Source, carbohydrate | 0.0650 | 0.0150 | 0.2050 | 0.0350 | 0.1300 | 0.0100 |
| PM1 | C11 | D-Melibiose | C-Source, carbohydrate | 0.0850 | 0.0150 | 0.2900 | 0.1500 | 0.1000 | 0.0100 |
| PM1 | C12 | Thymidine | C-Source, carbohydrate | 0.0950 | 0.0850 | 0.1600 | 0.0300 | 0.1150 | 0.0450 |
| PM1 | D01 | L-Asparagine | C-Source, amino acid | 0.1150 | 0.0350 | 0.1350 | 0.0050 | 0.1400 | 0.0100 |
| PM1 | D02 | D-Aspartic acid | C-Source, amino acid | 0.0750 | 0.0150 | 0.1350 | 0.0350 | 0.1400 | 0.0100 |
| PM1 | D03 | D-Glucosaminic acid | C-Source, carboxylic acid | 0.0550 | 0.0050 | 0.2650 | 0.1150 | 0.1350 | 0.0050 |
| PM1 | D04 | 1,2-Propanediol | C-Source, alcohol | 0.0650 | 0.0450 | 0.4600 | 0.1100 | 0.3650 | 0.0450 |
| PM1 | D05 | Tween 40 | C-Source, fatty acid | 21.2050 | 0.2250 | 17.2950 | 2.1050 | 30.7750 | 2.1250 |
| PM1 | D06 | a-Ketoglutaric acid | C-Source, carboxylic acid | 0.8200 | 0.1400 | 0.6050 | 0.1750 | 0.4600 | 0.1300 |
| PM1 | D07 | a-Ketobutyric acid | C-Source, carboxylic acid | 0.6150 | 0.0050 | 0.6600 | 0.0300 | 0.8550 | 0.0450 |
| PM1 | D08 | a-Methyl-D-Galactoside | C-Source, carbohydrate | 0.1350 | 0.0450 | 0.2950 | 0.1850 | 0.1500 | 0.0300 |
| PM1 | D09 | a-D-Lactose | C-Source, carbohydrate | 0.0850 | 0.0350 | 0.2100 | 0.0300 | 0.1200 | 0.0300 |
| PM1 | D10 | Lactulose | C-Source, carbohydrate | 0.0800 | 0.0400 | 0.2200 | 0.0400 | 0.1100 | 0.0300 |
| PM1 | D11 | Sucrose | C-Source, carbohydrate | 0.1200 | 0.0600 | 0.2650 | 0.0350 | 0.1700 | 0.0400 |
| PM1 | D12 | Uridine | C-Source, carbohydrate | 0.0650 | 0.0250 | 0.1200 | 0.0200 | 0.1000 | 0.0200 |
| PM1 | E01 | L-Glutamine | C-Source, amino acid | 0.2100 | 0.0800 | 0.2600 | 0.0200 | 0.1600 | 0.0300 |
| PM1 | E02 | m-Tartaric acid | C-Source, carboxylic acid | 0.0800 | 0.0300 | 0.1350 | 0.0550 | 0.1250 | 0.0150 |
| PM1 | E03 | D-Glucose-1-Phosphate | C-Source, carbohydrate | 0.0900 | 0.0400 | 0.1500 | 0.0200 | 0.1150 | 0.0150 |
| PM1 | E04 | D-Fructose-6-Phosphate | C-Source, carbohydrate | 0.1550 | 0.0050 | 0.2700 | 0.0100 | 0.2050 | 0.0050 |
| PM1 | E05 | Tween 80 | C-Source, fatty acid | 21.7700 | 0.4200 | 21.4750 | 0.4750 | 26.9400 | 0.3200 |
| PM1 | E06 | a-Hydroxyglutaric acid-g-La | C-Source, carboxylic acid | 0.0700 | 0.0100 | 0.1700 | 0.0400 | 0.1300 | 0.0100 |
| PM1 | E07 | a-Hydroxybutyric acid | C-Source, carboxylic acid | 0.3550 | 0.0450 | 0.2050 | 0.0450 | 1.2600 | 0.0100 |
| PM1 | E08 | b-Methyl-D-Glucoside | C-Source, carbohydrate | 0.2850 | 0.2450 | 0.3000 | 0.1600 | 0.5400 | 0.3900 |
| PM1 | E09 | Adonitol | C-Source, carbohydrate | 0.0750 | 0.0250 | 0.2000 | 0.0200 | 0.1300 | 0.0100 |
| PM1 | E10 | Maltotriose | C-Source, carbohydrate | 0.0200 | 0.0400 | 0.7500 | 0.0400 | 0.2100 | 0.0200 |
| PM1 | E11 | 2`-Deoxyadenosine | C-Source, carbohydrate | 0.0500 | 0.0300 | -0.0500 | 0.0300 | 0.0500 | 0.0300 |
| PM1 | E12 | Adenosine | C-Source, carbohydrate | 0.0650 | 0.0850 | 0.0450 | 0.0350 | 0.1300 | 0.0600 |
| PM1 | F01 | Gly-Asp | C-Source, amino acid | 0.0800 | 0.0300 | 0.2250 | 0.0550 | 0.1300 | 0.0100 |
| PM1 | F02 | Citric acid | C-Source, carboxylic acid | 0.0700 | 0.0200 | 0.1000 | 0.0000 | 0.1100 | 0.0200 |
| PM1 | F03 | m-Inositol | C-Source, carbohydrate | 0.0650 | 0.0150 | 0.1800 | 0.0200 | 0.1400 | 0.0100 |
| PM1 | F04 | D-Threonine | C-Source, amino acid | 0.0500 | 0.0000 | 0.1550 | 0.0150 | 0.1300 | 0.0300 |
| PM1 | F05 | Fumaric acid | C-Source, carboxylic acid | 5.8000 | 0.6400 | 7.2900 | 0.8100 | 5.1150 | 0.6350 |
| PM1 | F06 | Bromosuccinic acid | C-Source, carboxylic acid | 4.5050 | 0.1650 | 6.8000 | 0.2500 | 6.0900 | 0.4900 |
| PM1 | F07 | Propionic acid | C-Source, carboxylic acid | 6.0350 | 0.6050 | 9.6400 | 0.5000 | 16.2150 | 2.1150 |
| PM1 | F08 | Mucic acid | C-Source, carboxylic acid | 0.0500 | 0.0000 | 0.1350 | 0.0050 | 0.1700 | 0.0300 |
| PM1 | F09 | Glycolic acid | C-Source, carboxylic acid | 0.0350 | 0.0150 | 0.0700 | 0.0200 | 0.0400 | 0.0200 |
| PM1 | F10 | Glyoxylic acid | C-Source, carboxylic acid | 0.2250 | 0.0050 | 0.2400 | 0.0400 | 0.2450 | 0.0250 |
| PM1 | F11 | D-Cellobiose | C-Source, carbohydrate | 0.0600 | 0.0100 | 0.1850 | 0.0050 | 0.1400 | 0.0000 |
| PM1 | F12 | Inosine | C-Source, carbohydrate | 0.0200 | 0.0100 | 0.1400 | 0.0200 | 0.1250 | 0.0150 |
| PM1 | G01 | Gly-Glu | C-Source, amino acid | 0.1250 | 0.0350 | 0.1750 | 0.0750 | 0.1200 | 0.0300 |
| PM1 | G02 | Tricarballylic acid | C-Source, carboxylic acid | 0.1050 | 0.0150 | 0.1950 | 0.0250 | 0.1100 | 0.0500 |
| PM1 | G03 | L-Serine | C-Source, amino acid | 0.0650 | 0.0150 | 0.0750 | 0.0250 | 0.0600 | 0.0300 |
| PM1 | G04 | L-Threonine | C-Source, amino acid | 0.0650 | 0.0050 | 0.0950 | 0.0050 | 0.1250 | 0.0050 |
| PM1 | G05 | L-Alanine | C-Source, amino acid | 0.0250 | 0.0250 | 0.0550 | 0.0150 | 0.0650 | 0.0050 |
| PM1 | G06 | Ala-Gly | C-Source, amino acid | 0.0750 | 0.0350 | 0.1000 | 0.0500 | 0.3400 | 0.2000 |
| PM1 | G07 | Acetoacetic acid | C-Source, carboxylic acid | 1.6750 | 1.6050 | 1.1550 | 0.0050 | 1.3700 | 0.2300 |
| PM1 | G08 | N-Acetyl-D-Mannosamine | C-Source, carbohydrate | 0.0700 | 0.0200 | 0.1650 | 0.0250 | 0.1200 | 0.0400 |
| PM1 | G09 | Mono-Methylsuccinate | C-Source, carboxylic acid | 3.6650 | 0.1150 | 4.8800 | 0.1700 | 2.5500 | 2.3100 |
| PM1 | G10 | Methylpyruvate | C-Source, ester | 3.2650 | 0.0850 | 15.7250 | 0.6350 | 7.5700 | 3.1900 |
| PM1 | G11 | D-Malic acid | C-Source, carboxylic acid | 0.2050 | 0.0350 | 0.1850 | 0.0250 | 0.2350 | 0.0350 |
| PM1 | G12 | L-Malic acid | C-Source, carboxylic acid | 6.2450 | 0.2450 | 9.6000 | 0.0700 | 8.4300 | 0.1300 |
| PM1 | H01 | Gly-Pro | C-Source, amino acid | 0.0450 | 0.0050 | 0.1150 | 0.0550 | 0.0300 | 0.0100 |
| PM1 | H02 | p-Hydroxyphenyl Acetic acic | C-Source, carboxylic acid | 0.0900 | 0.0100 | 0.1150 | 0.0450 | 0.0900 | 0.0100 |
| PM1 | H03 | m-Hydroxyphenyl Acetic aci | C-Source, carboxylic acid | 6.1000 | 0.3800 | 9.1100 | 0.2000 | 10.8300 | 0.2900 |
| PM1 | H04 | Tyramine | C-Source, amine | 0.0500 | 0.0300 | 0.0400 | 0.0300 | 0.0500 | 0.0000 |
| PM1 | H05 | D-Psicose | C-Source, carbohydrate | 0.1700 | 0.0100 | 0.1750 | 0.0050 | 0.1750 | 0.0350 |
| PM1 | H06 | L-Lyxose | C-Source, carbohydrate | 17.4900 | 3.5000 | 17.9650 | 2.1450 | 13.2100 | 0.6500 |
| PM1 | H07 | Glucuronamide | C-Source, amide | 0.2200 | 0.0000 | 0.4350 | 0.1150 | 0.2350 | 0.0150 |
| PM1 | H08 | Pyruvic acid | C-Source, carboxylic acid | 2.5850 | 0.6950 | 16.9650 | 0.4950 | 9.6050 | 3.3950 |
| PM1 | H09 | L-Galactonic acid-g-Lactone | C-Source, carboxylic acid | 0.0850 | 0.0350 | 0.1650 | 0.0750 | 0.1950 | 0.0150 |
| PM1 | H10 | D-Galacturonic acid | C-Source, carboxylic acid | 0.0700 | 0.0300 | 0.1550 | 0.0350 | 0.1450 | 0.0350 |
| PM1 | H11 | Phenylethylamine | C-Source, amine | -0.0150 | 0.0450 | 0.1350 | 0.0050 | 0.0700 | 0.0000 |
| PM1 | H12 | 2-Aminoethanol | C-Source, alcohol | 0.0500 | 0.0100 | 0.0800 | 0.0300 | 0.0750 | 0.0350 |

| Plate | Well | Chemical | Mode of action | Maximmun growth rate | | | | | |
|---|---|---|---|---|---|---|---|---|---|
| | | | | PAM 2766 | | *R. equi* DSM 20307$^T$ | | *R. equi* 103S PAM 1126 | |
| | | | | Mean | SEM | Mean | SEM | Mean | SEM |
| PM2A | A01 | Negative Control | C-Source, negative contro | 0.1200 | 0.0300 | 0.1900 | 0.0300 | 0.2000 | 0.0100 |
| PM2A | A02 | Chondroitin Sulfate C | C-Source, polymer | 0.1250 | 0.0550 | 0.2100 | 0.1300 | 0.1900 | 0.0000 |
| PM2A | A03 | a-Cyclodextrin | C-Source, polymer | 0.1850 | 0.0350 | 0.7450 | 0.2850 | 0.9650 | 0.2350 |
| PM2A | A04 | b-Cyclodextrin | C-Source, polymer | 0.1150 | 0.0350 | 0.4450 | 0.1450 | 0.2600 | 0.0300 |
| PM2A | A05 | g-Cyclodextrin | C-Source, polymer | 0.2950 | 0.0550 | 0.7100 | 0.1100 | 0.8950 | 0.0550 |
| PM2A | A06 | Dextrin | C-Source, polymer | 0.2950 | 0.1150 | 0.7450 | 0.0950 | 0.2550 | 0.0150 |
| PM2A | A07 | Gelatin | C-Source, polymer | 0.1150 | 0.0450 | 0.7050 | 0.2150 | 0.1450 | 0.0450 |
| PM2A | A08 | Glycogen | C-Source, polymer | 0.0150 | 0.0350 | 0.4150 | 0.2550 | 0.1850 | 0.0550 |
| PM2A | A09 | Inulin | C-Source, polymer | 0.0550 | 0.0350 | 0.4550 | 0.3750 | 0.1700 | 0.1000 |
| PM2A | A10 | Laminarin | C-Source, polymer | 0.0900 | 0.0000 | 0.6550 | 0.2250 | 0.1350 | 0.0250 |
| PM2A | A11 | Mannan | C-Source, polymer | 0.0600 | 0.0000 | 0.2650 | 0.1350 | 0.1800 | 0.0200 |
| PM2A | A12 | Pectin | C-Source, polymer | 0.0600 | 0.0000 | 0.4200 | 0.0600 | 0.1300 | 0.0100 |
| PM2A | B01 | N-Acetyl-D-Galactosamine | C-Source, carbohydrate | 0.1000 | 0.0200 | 0.2800 | 0.0600 | 0.1850 | 0.0650 |
| PM2A | B02 | N-Acetyl-Neuraminic acid | C-Source, carboxylic acid | 0.0250 | 0.0150 | 0.1250 | 0.0450 | 0.0600 | 0.0000 |
| PM2A | B03 | b-D-Allose | C-Source, carbohydrate | 0.0950 | 0.0050 | 0.1450 | 0.0150 | 0.2000 | 0.0300 |
| PM2A | B04 | Amygdalin | C-Source, carbohydrate | 0.0300 | 0.0100 | 0.2700 | 0.1200 | 0.1250 | 0.0850 |
| PM2A | B05 | D-Arabinose | C-Source, carbohydrate | 0.6100 | 0.1600 | 1.3200 | 0.1700 | 1.2150 | 0.0150 |
| PM2A | B06 | D-Arabitol | C-Source, carbohydrate | 0.0550 | 0.0350 | 0.2950 | 0.1750 | 0.2450 | 0.1350 |
| PM2A | B07 | L-Arabitol | C-Source, carbohydrate | 0.0100 | 0.0200 | 0.2250 | 0.0850 | 0.1750 | 0.0750 |
| PM2A | B08 | Arbutin | C-Source, carbohydrate | 0.0300 | 0.0100 | 0.2500 | 0.1500 | 0.1300 | 0.0500 |
| PM2A | B09 | 2-Deoxy-D-Ribose | C-Source, carbohydrate | 8.6250 | 0.2050 | 11.9850 | 2.6350 | 9.0300 | 0.1400 |
| PM2A | B10 | i-Erythritol | C-Source, carbohydrate | 0.0150 | 0.0150 | 0.2500 | 0.1300 | 0.1400 | 0.0500 |
| PM2A | B11 | D-Fucose | C-Source, carbohydrate | 0.0550 | 0.0250 | 0.2800 | 0.1800 | 0.1450 | 0.0550 |
| PM2A | B12 | 3-0-b-D-Galactopyranosyl-D | C-Source, carbohydrate | 0.0350 | 0.0050 | 0.1250 | 0.1250 | 0.0450 | 0.0050 |
| PM2A | C01 | Gentiobiose | C-Source, carbohydrate | 0.0900 | 0.0200 | 0.3250 | 0.0550 | 0.2200 | 0.0300 |
| PM2A | C02 | L-Glucose | C-Source, carbohydrate | 0.0300 | 0.0100 | 0.2800 | 0.1500 | 0.1150 | 0.0450 |
| PM2A | C03 | D-Lactitol | C-Source, carbohydrate | 0.0100 | 0.0000 | 0.1600 | 0.0100 | 0.0950 | 0.0650 |
| PM2A | C04 | D-Melezitose | C-Source, carbohydrate | 0.0550 | 0.0050 | 0.5500 | 0.3200 | 0.2950 | 0.0950 |
| PM2A | C05 | Maltitol | C-Source, carbohydrate | 0.0100 | 0.0100 | 0.3050 | 0.1550 | 0.1400 | 0.0100 |
| PM2A | C06 | a-Methyl-D-Glucoside | C-Source, carbohydrate | 0.0400 | 0.0000 | 0.2750 | 0.1250 | 0.1150 | 0.0250 |
| PM2A | C07 | b-Methyl-D-Galactoside | C-Source, carbohydrate | 0.0250 | 0.0050 | 0.2650 | 0.1450 | 0.1250 | 0.0250 |
| PM2A | C08 | 3-Methylglucose | C-Source, carbohydrate | 0.0350 | 0.0050 | 0.1450 | 0.1250 | 0.1400 | 0.0300 |
| PM2A | C09 | b-Methyl-D-Glucuronic acid | C-Source, carboxylic acid | 0.0450 | 0.0150 | 0.1000 | 0.0100 | 0.0750 | 0.0150 |
| PM2A | C10 | a-Methyl-D-Mannoside | C-Source, carbohydrate | 0.0350 | 0.0150 | 0.1850 | 0.0650 | 0.1000 | 0.0100 |
| PM2A | C11 | b-Methyl-D-Xyloside | C-Source, carbohydrate | 0.0250 | 0.0150 | 0.1900 | 0.1500 | 0.0600 | 0.0200 |
| PM2A | C12 | Palatinose | C-Source, carbohydrate | 0.1700 | 0.0000 | 0.3250 | 0.1050 | 0.2350 | 0.0050 |
| PM2A | D01 | D-Raffinose | C-Source, carbohydrate | 0.1000 | 0.0100 | 0.1700 | 0.0100 | 0.2200 | 0.0200 |
| PM2A | D02 | Salicin | C-Source, carbohydrate | 0.0150 | 0.0150 | 0.1500 | 0.0500 | 0.0450 | 0.0450 |
| PM2A | D03 | Sedoheptulosan | C-Source, carbohydrate | 0.0600 | 0.0500 | 0.1600 | 0.0100 | 0.1050 | 0.0050 |
| PM2A | D04 | L-Sorbose | C-Source, carbohydrate | 0.0750 | 0.0150 | 0.1200 | 0.0100 | 0.4100 | 0.3000 |
| PM2A | D05 | Stachyose | C-Source, carbohydrate | 0.0400 | 0.0000 | 0.2000 | 0.0700 | 0.1800 | 0.0500 |
| PM2A | D06 | D-Tagatose | C-Source, carbohydrate | 0.2150 | 0.0150 | 0.3350 | 0.1250 | 0.2700 | 0.0600 |
| PM2A | D07 | Turanose | C-Source, carbohydrate | 0.0650 | 0.0050 | 0.5350 | 0.1450 | 0.3250 | 0.0350 |
| PM2A | D08 | Xylitol | C-Source, carbohydrate | 0.0200 | 0.0100 | 0.1550 | 0.0450 | 0.1300 | 0.0100 |
| PM2A | D09 | N-Acetyl-D-Glucosaminitol | C-Source, carbohydrate | 0.1200 | 0.0500 | 0.3100 | 0.1300 | 0.0950 | 0.0450 |
| PM2A | D10 | g-Amino-N-Butyric acid | C-Source, carboxylic acid | 0.0300 | 0.0000 | 0.2200 | 0.1600 | 0.0650 | 0.0050 |
| PM2A | D11 | d-Amino Valeric acid | C-Source, carboxylic acid | 0.0150 | 0.0050 | 0.1750 | 0.1250 | 0.0700 | 0.0400 |
| PM2A | D12 | Butyric acid | C-Source, carboxylic acid | 7.3450 | 0.1750 | 17.6150 | 0.1350 | 17.5300 | 1.3800 |
| PM2A | E01 | Capric acid | C-Source, carboxylic acid | 0.0850 | 0.0350 | 0.0750 | 0.0150 | 0.0500 | 0.0400 |
| PM2A | E02 | Caproic acid | C-Source, carboxylic acid | 2.2750 | 0.1950 | 18.7750 | 0.6750 | 15.6150 | 2.8350 |
| PM2A | E03 | Citraconic acid | C-Source, carboxylic acid | 0.0400 | 0.0300 | 0.1150 | 0.0350 | 0.1050 | 0.0450 |
| PM2A | E04 | D,L-Citramalic acid | C-Source, carboxylic acid | 0.0350 | 0.0050 | 0.1250 | 0.0150 | 0.1400 | 0.0300 |
| PM2A | E05 | D-Glucosamine | C-Source, carbohydrate | 0.8050 | 0.0050 | 0.8100 | 0.0100 | 0.7500 | 0.0300 |
| PM2A | E06 | 2-Hydroxybenzoic acid | C-Source, carboxylic acid | 0.0050 | 0.0150 | 0.0100 | 0.0400 | 0.0850 | 0.0350 |
| PM2A | E07 | 4-Hydroxybenzoic acid | C-Source, carboxylic acid | 0.0450 | 0.0150 | 6.9500 | 0.4100 | 6.8750 | 1.1850 |
| PM2A | E08 | b-Hydroxybutyric acid | C-Source, carboxylic acid | 0.5050 | 0.0150 | 5.3450 | 0.0650 | 5.2900 | 0.3000 |
| PM2A | E09 | g-Hydroxybutyric acid | C-Source, carboxylic acid | 0.0150 | 0.0050 | 0.0950 | 0.0350 | -0.0050 | 0.0150 |
| PM2A | E10 | a-Keto-Valeric acid | C-Source, carboxylic acid | 1.3400 | 0.0100 | 3.0900 | 0.6200 | 1.8100 | 0.6900 |
| PM2A | E11 | Itaconic acid | C-Source, carboxylic acid | 0.0050 | 0.0150 | 0.2050 | 0.1450 | 0.0500 | 0.0200 |
| PM2A | E12 | 5-Keto-D-Gluconic acid | C-Source, carboxylic acid | 1.0700 | 0.1700 | 1.2550 | 0.0350 | 1.3500 | 0.1600 |
| PM2A | F01 | D-Lactic acid Methyl Ester | C-Source, ester | 1.0850 | 0.2650 | 5.7400 | 0.8800 | 0.1850 | 0.0950 |
| PM2A | F02 | Malonic acid | C-Source, carboxylic acid | 0.0500 | 0.0300 | 0.1350 | 0.0050 | 0.1050 | 0.0550 |
| PM2A | F03 | Melibionic acid | C-Source, carbohydrate | 0.0400 | 0.0400 | 0.0550 | 0.1050 | 0.1250 | 0.0250 |
| PM2A | F04 | Oxalic acid | C-Source, carboxylic acid | 0.0300 | 0.0000 | 0.1100 | 0.0300 | 0.1300 | 0.0000 |
| PM2A | F05 | Oxalomalic acid | C-Source, carboxylic acid | 0.9200 | 0.2500 | 1.2450 | 0.0550 | 1.3050 | 0.1050 |
| PM2A | F06 | Quinic acid | C-Source, carboxylic acid | 0.0700 | 0.0200 | 0.2550 | 0.1250 | 0.1550 | 0.0050 |
| PM2A | F07 | D-Ribono-1,4-Lactone | C-Source, carboxylic acid | 0.0800 | 0.0100 | 0.0900 | 0.0100 | 0.1050 | 0.0050 |
| PM2A | F08 | Sebacic acid | C-Source, carboxylic acid | 3.3450 | 0.3050 | 0.4950 | 0.0650 | 0.5100 | 0.0300 |
| PM2A | F09 | Sorbic acid | C-Source, carboxylic acid | 2.2300 | 0.2300 | 3.1400 | 1.0500 | 3.4700 | 1.0100 |
| PM2A | F10 | Succinamic acid | C-Source, carboxylic acid | 0.2000 | 0.0800 | 2.7000 | 0.2400 | 0.1250 | 0.0050 |
| PM2A | F11 | D-Tartaric acid | C-Source, carboxylic acid | 0.0450 | 0.0150 | 0.0800 | 0.0200 | 0.0950 | 0.0250 |
| PM2A | F12 | L-Tartaric acid | C-Source, carboxylic acid | 0.0000 | 0.0300 | 0.1400 | 0.0100 | 0.0950 | 0.0050 |
| PM2A | G01 | Acetamide | C-Source, amide | 0.0850 | 0.0450 | 0.1350 | 0.0750 | 6.7900 | 6.6500 |
| PM2A | G02 | L-Alaninamide | C-Source, amide | 0.0150 | 0.0250 | 0.0450 | 0.0250 | 0.0700 | 0.0300 |
| PM2A | G03 | N-Acetyl-L-Glutamic acid | C-Source, amino acid | 0.0250 | 0.0050 | 0.1200 | 0.0500 | 0.1150 | 0.0150 |
| PM2A | G04 | L-Arginine | C-Source, amino acid | 0.0150 | 0.0050 | 0.0850 | 0.0050 | 0.0850 | 0.0050 |
| PM2A | G05 | Glycine | C-Source, amino acid | 0.0350 | 0.0150 | 0.0550 | 0.0050 | 0.0800 | 0.0000 |
| PM2A | G06 | L-Histidine | C-Source, amino acid | 4.7400 | 0.1200 | 1.7250 | 1.4650 | 4.0450 | 0.2750 |
| PM2A | G07 | L-Homoserine | C-Source, amino acid | 0.0200 | 0.0000 | 0.1000 | 0.0300 | 0.0950 | 0.0050 |
| PM2A | G08 | Hydroxy-L-Proline | C-Source, amino acid | 0.0450 | 0.0350 | 0.1300 | 0.0100 | 0.1200 | 0.0100 |
| PM2A | G09 | L-Isoleucine | C-Source, amino acid | 1.5050 | 1.4750 | 0.1400 | 0.0200 | 0.1700 | 0.0800 |
| PM2A | G10 | L-Leucine | C-Source, amino acid | 3.1350 | 3.0150 | 0.2000 | 0.0300 | 0.0950 | 0.0050 |
| PM2A | G11 | L-Lysine | C-Source, amino acid | 0.0050 | 0.0050 | 0.1550 | 0.0950 | 0.0250 | 0.0050 |
| PM2A | G12 | L-Methionine | C-Source, amino acid | 0.0450 | 0.0550 | 0.0950 | 0.0150 | 0.1350 | 0.0950 |
| PM2A | H01 | L-Ornithine | C-Source, amino acid | 0.0550 | 0.0450 | 0.0950 | 0.0050 | 0.0800 | 0.0500 |
| PM2A | H02 | L-Phenylalanine | C-Source, amino acid | 0.0700 | 0.0100 | 0.0700 | 0.0100 | 0.0800 | 0.0500 |
| PM2A | H03 | L-Pyroglutamic acid | C-Source, amino acid | 0.0150 | 0.0250 | 0.1650 | 0.0250 | 0.1150 | 0.0150 |
| PM2A | H04 | L-Valine | C-Source, amino acid | 0.0650 | 0.0150 | 0.0900 | 0.0400 | 0.1150 | 0.0250 |
| PM2A | H05 | D,L-Carnitine | C-Source, carboxylic acid | 0.0550 | 0.0450 | 0.0950 | 0.0250 | 0.0850 | 0.0050 |
| PM2A | H06 | sec-Butylamine | C-Source, amine | 0.0450 | 0.0350 | 0.1050 | 0.0150 | 0.0850 | 0.0150 |
| PM2A | H07 | D,L-Octopamine | C-Source, amine | 0.0550 | 0.0350 | 0.1400 | 0.0000 | 0.1100 | 0.0000 |
| PM2A | H08 | Putrescine | C-Source, amine | 0.0550 | 0.0050 | 0.0850 | 0.0150 | 0.0600 | 0.0100 |
| PM2A | H09 | Dihydroxyacetone | C-Source, alcohol | 3.8400 | 0.1000 | 9.0450 | 5.3250 | 3.6750 | 0.0150 |
| PM2A | H10 | 2,3-Butanediol | C-Source, alcohol | 0.0300 | 0.0100 | 0.1100 | 0.0100 | 0.0750 | 0.0150 |
| PM2A | H11 | 2,3-Butanone | C-Source, alcohol | 0.5300 | 0.0600 | 0.5400 | 0.0000 | 0.9150 | 0.1250 |
| PM2A | H12 | 3-Hydroxy-2-butanone | C-Source, alcohol | 0.0300 | 0.0400 | 0.0600 | 0.0100 | 0.0300 | 0.0100 |

| Plate | Well | Chemical | Mode of action | Maximmun growth rate | | | | | |
|---|---|---|---|---|---|---|---|---|---|
| | | | | PAM 2766 | | *R. equi* DSM 20307$^T$ | | *R. equi* 103S PAM 1126 | |
| | | | | Mean | SEM | Mean | SEM | Mean | SEM |
| PM3B | A01 | Negative Control | N-Source, Negative contrc | 1.1900 | 0.0141 | 0.9050 | 0.1485 | 1.4350 | 0.0495 |
| PM3B | A02 | Ammonia | N-Source, inorganic | 9.4900 | 0.0707 | 12.4950 | 1.7466 | 21.5500 | 2.0365 |
| PM3B | A03 | Nitrite | N-Source, inorganic | 6.2150 | 0.2333 | 8.0900 | 0.4525 | 13.7350 | 0.0071 |
| PM3B | A04 | Nitrate | N-Source, inorganic | 5.8650 | 0.2192 | 6.9650 | 0.0778 | 13.3800 | 0.0849 |
| PM3B | A05 | Urea | N-Source, other | 7.7400 | 0.1980 | 8.1550 | 0.4031 | 16.2800 | 2.1496 |
| PM3B | A06 | Biuret | N-Source, other | 1.7950 | 0.1768 | 0.6600 | 0.0141 | 1.8500 | 0.0424 |
| PM3B | A07 | L-Alanine | N-Source, amino acid | 1.9450 | 0.0919 | 2.1700 | 0.0424 | 2.2350 | 0.0636 |
| PM3B | A08 | L-Arginine | N-Source, amino acid | 1.4600 | 0.0707 | 0.7050 | 0.1344 | 1.5550 | 0.2475 |
| PM3B | A09 | L-Asparagine | N-Source, amino acid | 5.3200 | 0.0000 | 10.9250 | 0.6435 | 12.0250 | 0.2475 |
| PM3B | A10 | L-Aspartic acid | N-Source, amino acid | 6.1350 | 1.3364 | 7.1200 | 0.4667 | 7.0700 | 0.4950 |
| PM3B | A11 | L-Cysteine | N-Source, amino acid | 0.7050 | 0.0778 | 4.6500 | 0.3536 | 5.6750 | 0.1626 |
| PM3B | A12 | L-Glutamic acid | N-Source, amino acid | 5.2800 | 0.6223 | 9.9650 | 0.2333 | 7.3800 | 0.7495 |
| PM3B | B01 | L-Glutamine | N-Source, amino acid | 9.5900 | 0.3111 | 14.7000 | 0.6081 | 21.5650 | 0.3748 |
| PM3B | B02 | Glycine | N-Source, amino acid | 1.7350 | 0.2192 | 2.1850 | 0.0212 | 3.1250 | 0.0354 |
| PM3B | B03 | L-Histidine | N-Source, amino acid | 10.9450 | 0.3465 | 13.3550 | 0.4455 | 18.6800 | 1.3435 |
| PM3B | B04 | L-Isoleucine | N-Source, amino acid | 2.9600 | 0.3960 | 1.3850 | 0.0354 | 3.6600 | 0.4808 |
| PM3B | B05 | L-Leucine | N-Source, amino acid | 4.3900 | 0.1414 | 9.6950 | 0.4172 | 9.9050 | 0.1909 |
| PM3B | B06 | L-Lysine | N-Source, amino acid | 0.9650 | 0.3041 | 8.8200 | 0.7495 | 10.7550 | 0.0919 |
| PM3B | B07 | L-Methionine | N-Source, amino acid | 0.8600 | 0.2970 | 0.5750 | 0.1202 | 1.2750 | 0.2192 |
| PM3B | B08 | L-Phenylalanine | N-Source, amino acid | 4.3300 | 0.0849 | 7.6600 | 1.0465 | 9.1750 | 0.2051 |
| PM3B | B09 | L-Proline | N-Source, amino acid | 1.2050 | 0.0495 | 0.8900 | 0.4950 | 1.2250 | 0.0212 |
| PM3B | B10 | L-Serine | N-Source, amino acid | 0.4350 | 0.0778 | 0.5400 | 0.0707 | 0.9850 | 0.0212 |
| PM3B | B11 | L-Threonine | N-Source, amino acid | 0.9850 | 0.0636 | 0.5000 | 0.1131 | 1.8400 | 0.4384 |
| PM3B | B12 | L-Tryptophan | N-Source, amino acid | 3.9550 | 0.9546 | 9.6800 | 0.2263 | 9.7400 | 0.3111 |
| PM3B | C01 | L-Tyrosine | N-Source, amino acid | 2.0050 | 0.1485 | 6.7450 | 1.1809 | 8.1800 | 0.1556 |
| PM3B | C02 | L-Valine | N-Source, amino acid | 2.6800 | 0.1273 | 1.6400 | 0.3818 | 3.2050 | 0.0354 |
| PM3B | C03 | D-Alanine | N-Source, amino acid | 2.3150 | 0.1202 | 2.5050 | 0.0071 | 3.0550 | 0.1485 |
| PM3B | C04 | D-Asparagine | N-Source, amino acid | 2.3050 | 0.2051 | 1.1150 | 0.2475 | 2.6500 | 0.0283 |
| PM3B | C05 | D-Aspartic acid | N-Source, amino acid | 2.5650 | 0.3606 | 1.0350 | 0.3041 | 2.6050 | 0.1626 |
| PM3B | C06 | D-Glutamic acid | N-Source, amino acid | 1.4700 | 0.0283 | 0.7500 | 0.0283 | 1.7850 | 0.3889 |
| PM3B | C07 | D-Lysine | N-Source, amino acid | 4.1600 | 0.0990 | 7.9000 | 0.5798 | 13.2550 | 0.3889 |
| PM3B | C08 | D-Serine | N-Source, amino acid | 1.5300 | 0.0849 | 1.0200 | 0.3253 | 1.5950 | 0.0495 |
| PM3B | C09 | D-Valine | N-Source, amino acid | 1.4350 | 0.2758 | 0.6200 | 0.0283 | 1.5050 | 0.0636 |
| PM3B | C10 | L-Citrulline | N-Source, amino acid | 1.9800 | 0.4950 | 8.8800 | 0.1980 | 6.6850 | 0.1626 |
| PM3B | C11 | L-Homoserine | N-Source, amino acid | 1.7900 | 0.0990 | 1.2300 | 0.2404 | 2.5250 | 0.2051 |
| PM3B | C12 | L-Ornithine | N-Source, amino acid | 0.9500 | 0.0424 | 6.2250 | 0.1202 | 10.7650 | 0.0495 |
| PM3B | D01 | N-Acetyl-L-Glutamic acid | N-Source, amino acid | 1.0500 | 0.1414 | 0.7550 | 0.1626 | 1.3500 | 0.1131 |
| PM3B | D02 | N-Phthaloyl-L-Glutamic acid | N-Source, amino acid | 1.3250 | 0.0495 | 0.5600 | 0.0849 | 1.5850 | 0.0636 |
| PM3B | D03 | L-Pyroglutamic acid | N-Source, amino acid | 1.8600 | 0.2828 | 0.8550 | 0.0212 | 2.4800 | 0.0141 |
| PM3B | D04 | Hydroxylamine | N-Source, other | 0.2100 | 0.0424 | 0.1600 | 0.0000 | 0.5500 | 0.5657 |
| PM3B | D05 | Methylamine | N-Source, other | 2.4250 | 0.1485 | 5.5700 | 0.1838 | 12.8100 | 1.1031 |
| PM3B | D06 | N-Amylamine | N-Source, other | 5.3100 | 0.3536 | 3.2650 | 0.2192 | 8.5700 | 0.2263 |
| PM3B | D07 | N-Butylamine | N-Source, other | 5.3600 | 0.1697 | 6.0150 | 0.5869 | 9.3600 | 0.0990 |
| PM3B | D08 | Ethylamine | N-Source, other | 4.4850 | 0.4031 | 5.9900 | 0.3818 | 10.4950 | 0.0212 |
| PM3B | D09 | Ethanolamine | N-Source, other | 4.5300 | 0.0990 | 5.6150 | 0.2192 | 9.8300 | 0.2121 |
| PM3B | D10 | Ethylenediamine | N-Source, other | 0.3300 | 0.0566 | 0.3600 | 0.0566 | 0.8700 | 0.6788 |
| PM3B | D11 | Putrescine | N-Source, other | 6.7200 | 0.5940 | 9.6900 | 0.1697 | 11.2250 | 0.3889 |
| PM3B | D12 | Agmatine | N-Source, other | 2.1900 | 0.7637 | 4.2900 | 0.0707 | 2.7500 | 0.0424 |
| PM3B | E01 | Histamine | N-Source, other | 4.8000 | 0.2687 | 6.3500 | 0.0424 | 13.4200 | 0.1414 |
| PM3B | E02 | b-Phenylethylamine | N-Source, other | 3.3050 | 0.9405 | 0.8400 | 0.1697 | 8.1300 | 0.0424 |
| PM3B | E03 | Tyramine | N-Source, other | 3.9900 | 1.0465 | 4.5700 | 1.1738 | 5.5400 | 0.2970 |
| PM3B | E04 | Acetamide | N-Source, other | 4.3950 | 0.0636 | 9.4250 | 0.1485 | 21.0850 | 1.0677 |
| PM3B | E05 | Formamide | N-Source, other | 9.4750 | 0.4313 | 9.3900 | 0.0707 | 17.8400 | 1.3718 |
| PM3B | E06 | Glucuronamide | N-Source, other | 8.0950 | 0.2758 | 7.1550 | 0.4313 | 17.3200 | 0.0424 |
| PM3B | E07 | D,L-Lactamide | N-Source, other | 8.1400 | 1.3718 | 7.6200 | 0.3677 | 8.4600 | 0.4384 |
| PM3B | E08 | D-Glucosamine | N-Source, other | 0.3850 | 0.0495 | 0.3250 | 0.0212 | 2.2050 | 2.7365 |
| PM3B | E09 | D-Galactosamine | N-Source, other | 9.1050 | 0.1061 | 4.4950 | 0.0495 | 12.7250 | 0.0071 |
| PM3B | E10 | D-Mannosamine | N-Source, other | 1.6450 | 0.0919 | 1.0500 | 0.0141 | 13.6450 | 0.8839 |
| PM3B | E11 | N-Acetyl-D-Glucosamine | N-Source, other | 1.2050 | 0.1202 | 0.5250 | 0.1626 | 1.2000 | 0.1697 |
| PM3B | E12 | N-Acetyl-D-Galactosamine | N-Source, other | 0.9150 | 0.1909 | 0.4200 | 0.0707 | 0.8800 | 0.1131 |
| PM3B | F01 | N-Acetyl-D-Mannosamine | N-Source, other | 0.9850 | 0.0071 | 0.5400 | 0.0283 | 1.2250 | 0.0071 |
| PM3B | F02 | Adenine | N-Source, other | 7.2600 | 0.3111 | 5.7950 | 0.5869 | 8.7800 | 0.8627 |
| PM3B | F03 | Adenosine | N-Source, other | 4.4650 | 0.9405 | 0.1850 | 0.0212 | 3.1050 | 1.2092 |
| PM3B | F04 | Cytidine | N-Source, other | 2.0350 | 0.2192 | 1.1250 | 0.1909 | 2.9900 | 0.4525 |
| PM3B | F05 | Cytosine | N-Source, other | 7.6550 | 0.0071 | 7.9050 | 0.2475 | 10.1500 | 0.8061 |
| PM3B | F06 | Guanine | N-Source, other | 4.0500 | 4.3841 | 0.9450 | 0.0495 | 12.0400 | 14.7502 |
| PM3B | F07 | Guanosine | N-Source, other | 0.5950 | 0.0071 | 0.5700 | 0.0141 | 1.0900 | 0.0849 |
| PM3B | F08 | Thymine | N-Source, other | 2.1250 | 0.1909 | 8.6300 | 0.8344 | 14.3700 | 0.2687 |
| PM3B | F09 | Thymidine | N-Source, other | 1.3400 | 0.0283 | 0.6600 | 0.1556 | 1.5200 | 0.0283 |
| PM3B | F10 | Uracil | N-Source, other | 1.3700 | 0.0424 | 0.7050 | 0.0495 | 1.6450 | 0.4596 |
| PM3B | F11 | Uridine | N-Source, other | 0.8950 | 0.0071 | 0.3950 | 0.0778 | 1.1150 | 0.0919 |
| PM3B | F12 | Inosine | N-Source, other | 1.0400 | 0.1131 | 0.3800 | 0.0283 | 0.9550 | 0.0071 |
| PM3B | G01 | Xanthine | N-Source, other | 0.0550 | 0.0354 | 0.2900 | 0.2687 | 1.2800 | 0.4525 |
| PM3B | G02 | Xanthosine | N-Source, other | 1.2350 | 0.0354 | 0.6550 | 0.0636 | 1.4050 | 0.0354 |
| PM3B | G03 | Uric acid | N-Source, other | 7.1300 | 0.5657 | 6.7900 | 0.4101 | 7.0650 | 0.3606 |
| PM3B | G04 | Alloxan | N-Source, other | 3.9900 | 0.0849 | 3.5300 | 0.1980 | 3.7700 | 0.1273 |
| PM3B | G05 | Allantoin | N-Source, other | 5.3550 | 0.2475 | 3.5600 | 0.6647 | 5.2950 | 0.1909 |
| PM3B | G06 | Parabanic acid | N-Source, other | 6.5200 | 0.0707 | 7.9300 | 0.2121 | 12.3300 | 0.0283 |
| PM3B | G07 | D,L-a-Amino-N-Butyric acid | N-Source, other | 4.5600 | 0.8910 | 1.3400 | 0.7495 | 5.2150 | 0.1485 |
| PM3B | G08 | g-Amino-N-Butyric acid | N-Source, other | 1.1550 | 0.1061 | 1.2450 | 0.0071 | 1.7500 | 0.3818 |
| PM3B | G09 | e-Amino-N-Caproic acid | N-Source, other | 1.2150 | 0.0212 | 6.0100 | 0.4525 | 11.7900 | 0.2546 |
| PM3B | G10 | D,L-a-Amino-Caprylic acid | N-Source, other | 0.5150 | 0.0636 | 0.2700 | 0.0990 | 0.5450 | 0.5445 |
| PM3B | G11 | d-Amino-N-Valeric acid | N-Source, other | 0.8500 | 0.0566 | 6.0950 | 0.6435 | 9.7800 | 0.8627 |
| PM3B | G12 | a-Amino-N-Valeric acid | N-Source, other | 3.1050 | 1.0112 | 6.7900 | 0.4384 | 6.3300 | 0.6930 |
| PM3B | H01 | Ala-Asp | N-Source, peptide | 1.6900 | 0.1414 | 0.3950 | 0.0212 | 0.8300 | 0.0141 |
| PM3B | H02 | Ala-Gln | N-Source, peptide | 8.0350 | 0.1061 | 10.0550 | 0.9970 | 6.1300 | 0.0990 |
| PM3B | H03 | Ala-Glu | N-Source, peptide | 3.7000 | 0.0000 | 0.8500 | 0.0566 | 1.2850 | 0.1909 |
| PM3B | H04 | Ala-Gly | N-Source, peptide | 2.4300 | 0.0707 | 3.5850 | 0.3465 | 2.4900 | 0.1980 |
| PM3B | H05 | Ala-His | N-Source, peptide | 6.5650 | 0.6435 | 6.8850 | 0.7990 | 4.2550 | 0.5586 |
| PM3B | H06 | Ala-Leu | N-Source, peptide | 4.1450 | 0.1202 | 4.2900 | 0.3536 | 3.9400 | 0.4525 |
| PM3B | H07 | Ala-Thr | N-Source, peptide | 1.4750 | 0.1202 | 0.3850 | 0.0495 | 3.2350 | 0.1626 |
| PM3B | H08 | Gly-Asn | N-Source, peptide | 2.4750 | 0.0071 | 0.9700 | 0.0990 | 1.7600 | 0.2404 |
| PM3B | H09 | Gly-Gln | N-Source, peptide | 4.2250 | 0.0495 | 5.5100 | 0.4101 | 2.7350 | 0.1768 |
| PM3B | H10 | Gly-Glu | N-Source, peptide | 2.0700 | 0.0566 | 0.4400 | 0.0000 | 0.7150 | 0.0495 |
| PM3B | H11 | Gly-Met | N-Source, peptide | 2.4350 | 0.3323 | 3.6400 | 0.1838 | 3.9150 | 0.0212 |
| PM3B | H12 | Met-Ala | N-Source, peptide | 1.9150 | 0.0354 | 3.0350 | 0.1626 | 3.6700 | 0.0707 |
